# A key regulator of missing-self innate immunity is polymorphic and under diversifying selection

**DOI:** 10.64898/2025.12.10.693425

**Authors:** Rocco F. Notarnicola, Magdalena Herdegen-Radwan, Joanna Różańska-Wróbel, Mateusz Konczal, Karolina Przesmycka, Petr Kotlík, Wiesław Babik, Jacek Radwan

## Abstract

Host-parasite co-evolution drives the diversification of host immune genes involved in the recognition of pathogen antigens and molecular patterns. In contrast, the immune genes involved in self-recognition and inhibition of immune responses against self-cells (missing-self immunity) are expected to be evolutionarily constrained. However, many pathogens, such as the Lyme disease agent *Borrelia*, hijack these genes to evade the immune system and may therefore select for their diversification. How these contrasting but concurrent selective forces shape the evolution of missing-self regulators is not clearly understood. To fill this gap, we investigated polymorphism and molecular signatures of selection acting on a missing-self regulator, the Complement Factor H (CFH), in bank vole populations, which are an important wild reservoir for *Borrelia*. We then compared the geographic structuring in the CFH domain interacting with *Borrelia* (CCP 20) against a genomic background represented by RAD-seq markers. We found signals of positive and diversifying selection at CCP 20, suggesting that CFH evolved in response to pressures from pathogens. Additionally, we found other innate immunity genes within the alternative complement pathway, which is regulated by CFH, under diversifying selection, highlighting its involvement in host-parasite coevolution. This study demonstrates that an innate missing-self sensor in a wild vertebrate is under diversifying selection, likely driven by pathogens.

## Introduction

Immune genes are often found to evolve at faster rates than the genomic average, likely in response to ongoing adaptation of ever-evolving pathogens (Obbard et al., 2009; Enard et al., 2016; Shultz and Sackton, 2019). Host immune genes coding for effectors interacting with parasites, such as the Major Histocompatibility Complex (MHC) of the adaptive immunity, or the Pattern Recognition Receptors (PRR) of the innate immunity, are important components of these dynamics. Indeed, MHC genes encoding proteins crucial in self/non-self-recognition, initiating the adaptive immune response, have become a paradigm for host-pathogen coevolution. They are characterised by high amino acid substitution rates at regions involved in binding to antigens (Hughes and Nei, 1988, 1989; Garrigan and Hedrick, 2003) and extreme polymorphism, likely maintained by balancing selection imposed by parasites (Apanius et al., 1997; Radwan et al., 2020; Migalska et al., 2022). Selection pressures on innate immunity genes have been less studied, but the evidence is accumulating for fast molecular evolution in the regions involved in interactions with pathogens (Wlasiuk et al., 2009; Tschirren et al., 2013; Tian et al., 2018; Lundberg et al., 2020; Nandakumar et al., 2023). These studies mainly investigated genes encoding proteins that recognise pathogen-associated molecular patterns (PAMPs), while research exploring the evolution of innate immunity genes that mediate the missing-self signal is scant.

Missing identity controls the activity of the alternative complement pathway, an ancient system shared by vertebrates with ascidians and cephalochordates (Nonaka and Kimura, 2006). The complement system consists of multiple proteins involved in the phagocytosis and clearance of pathogens and damaged cells, the induction of inflammation, and the regulation of innate and adaptive immunity (Reis et al., 2019). Unlike the classical and lectin pathways, the alternative pathway is in a constant low-level activation state, and destruction of host healthy cells is prevented by protein regulators (Reis et al., 2019). One such regulator is the Complement Factor H (CFH), a 155 kDa protein with conserved structure and function (Schmidt et al., 2008; Pouw et al., 2015). The CFH gene consists of 22 exons that form 20 Complement Control Protein (CCP) domains, also known as Short Consensus Repeats or *sushi*. The 20 domains contain several glycosylation sites and binding sites to polyanions and to complement Component 3b (C3b; Schmidt et al., 2008; Pouw et al., 2015). The binding of CFH leads to C3b inactivation and/or prevents the formation of the C3 convertase (Meri et al., 2013; Schmidt et al., 2016), a protein that initiates the complement cascade that results in the opsonisation and destruction of pathogens. Host cells are protected from complement activation via binding of CFH to polyanions (sialic acids, glycosaminoglycans, phospholipids), molecules present on vertebrate cells but not on microorganisms (Meri, 2016) - therefore, bound CFH represents a self-identity signal. However, some pathogens, including *Borrelia burgdorferi*, *Pseudomonas aeruginosa*, and *Haemophilus influenzae,* evolved surface proteins that bind to CFH and thus hijack this self-identity signal (Meri et al., 2013). Although CFH can be expected to be conserved, being an immune regulator, hijacking by pathogens may trigger adaptive evolution of its sequence to counter this effect (Carrillo-Bustamante et al. 2016). However, how CFH evolves in response to pathogens is not well understood.

Many of the bacterial proteins that hijack CFH to mimic self and evade complement activation, including *Borrelia*’s outer surface protein E (OspE), bind to CCP domain 20 (Bhattacharjee et al. 2013; Meri et al. 2013), which also contains polyanion binding sites (Meri, 2016) and may therefore be subject to complex evolutionary processes/pressures. Cagliani et al. (2016) analysed molecular evolution of the complement system among primates and identified signals of positive selection in regions, such as CCP 6-7 and 19-20 of CFH, involved in interactions with pathogens belonging to different phyla. Furthermore, using *in-silico* structural modelling, the authors showed that *Borrelia*’s OspE residues interacting with CCP 20 were themselves under positive selection. In captive rhesus macaques, intraspecific variation within CCP 6 was linked to the binding efficiency of a meningococcal protein that evades immune recognition (Konar et al., 2015). However, the relevance of these interactions to the evolution of CFH is unclear, given that there is no evidence that *Borrelia* can naturally infect primates (Wolcott et al. 2021), while humans are only incidental and dead-end hosts, from which transmission to other competent hosts is interrupted. Additionally, in humans, mutations in CCP 19-20 were linked to several autoimmune diseases, such as age-related macular degeneration (Rodriguez et al., 2014; Ferluga et al., 2017), highlighting possible constraints on CFH evolution imposed by its involvement in self-recognition. Whether such constraints restrict polymorphism at CFH is unknown, as, to our knowledge, no study systematically investigated CFH polymorphism in any wild populations. Overall, it remains unclear how the contrasting selective pressures, exerted by pathogens and the role in self-recognition, shape CFH evolution.

Here, we characterise CFH polymorphism and analyse signatures of selection in wild populations of bank voles (*Clethrionomys glareolus*), a small woodland rodent with widespread distribution in Europe (Kotlik et al., 2006). After the Last Glacial Maximum (LGM), *C. glareolus* colonised Europe from several refugia, which included Carpathian and Eastern refugia, giving rise to contemporary populations that came back into contact in central Poland (Wójcik et al., 2010; Marková et al., 2020). A third lineage (Western) was also identified in previous studies, though it contributes some ancestry primarily in southern Poland (Wójcik et al., 2010; Marková et al., 2020). The analysis of genetic differentiation between two populations representing these two post-glacial colonisation areas, based on heart transcriptomes, revealed CFH as one of the most divergent genes between the two lineages (d_xy_ = 0.017; max F_STnonsyn_ = 1; mean F_ST_ = 0.72; Niedziałkowska et al., 2023; Figure S1). This polymorphism may be functionally relevant given that the bank vole is one of the main reservoirs for *Borrelia burgdorferi sensu lato* in Europe (Humair et al., 1999; van Duijvendijk et al., 2015). However, the analyses in Niedziałkowska et al. (2023) were based only on 18 individuals from two populations (one population per post-glacial colonisation area), therefore their results may represent an effect of sampling rather than regional structuring. Furthermore, no formal tests for signals of selection acting on the polymorphic sites in CFH were conducted.

In this study, we analyse CFH polymorphism within and across 13 bank vole populations spanning the two major post-glacial colonisation areas (Eastern and Carpathian). We implement two approaches to understand the nature of selection shaping polymorphism in this gene. The first one relies on the relative rate of non-synonymous to synonymous substitutions, which can detect positive selection over the history of the species (Garrigan and Hedrick, 2003). The second approach investigates how contemporary selection shapes the differentiation of allele frequencies between populations. Loci under balancing selection tend to remain polymorphic within populations, resulting in lower differentiation of allele frequencies between populations (Schierup et al., 2000). In contrast, local adaptation leads to stronger population structure and higher allele frequency differentiation in loci under selection compared to the genomic average. For this second approach, we conducted robust outlier tests with BayPass, which factors in complex demographic histories (shared ancestry and gene flow) between populations (Günther and Coop, 2013; Gautier, 2015), and compared the extent/levels of CFH differentiation across populations with genome-wide differentiation assessed by RADseq markers. Additionally, we checked whether the genes in proximity to outlier RAD markers were involved with immunity, particularly with the complement pathway.

## Results

### CFH is polymorphic and under positive selection, particularly at the CCP 20 domain

Amplification and ONT sequencing of full-length transcripts from 19 individuals revealed 22 unique full-length CFH sequences, with an expected length (3,748-3,769 bp) congruent with the well-annotated gene in *Rattus norvegicus* (3,708 bp; NM_130409.2) and *Mus musculus* (3,702 bp; BC066092-1). The analysis of CFH sequences may be challenging because of the presence of similar paralogs (CFH-related genes) and pseudogenes. Therefore, we further confirmed that the sequences represented a single gene by aligning them to a database of expressed CFH obtained by amplifying cDNA of CCP 17-20 from a subsample of 50 individuals (*see Methods*). CFH-related genes lack CCP 17 and/or 18; thus, our cDNA sequences were indeed CFH (Pouw et al., 2015). Phylogenetic analysis of full-length CFH revealed three clades (Figure 1). The average sequence divergence between variants was 0.014. The most polymorphic domain was CCP 20 (189 bp), containing 20 SNPs, 15 of which were non-synonymous (Figure 2). A re-analysis of data from Niedziałkowska et al. (2023), following the method of Nei and Gojobori (1986) to calculate divergence, showed that CFH is among the 1.47% most diverged transcripts for non-synonymous SNPs and 0.96% for synonymous SNPs between two populations in NE and Central Poland.

**Figure 1:**
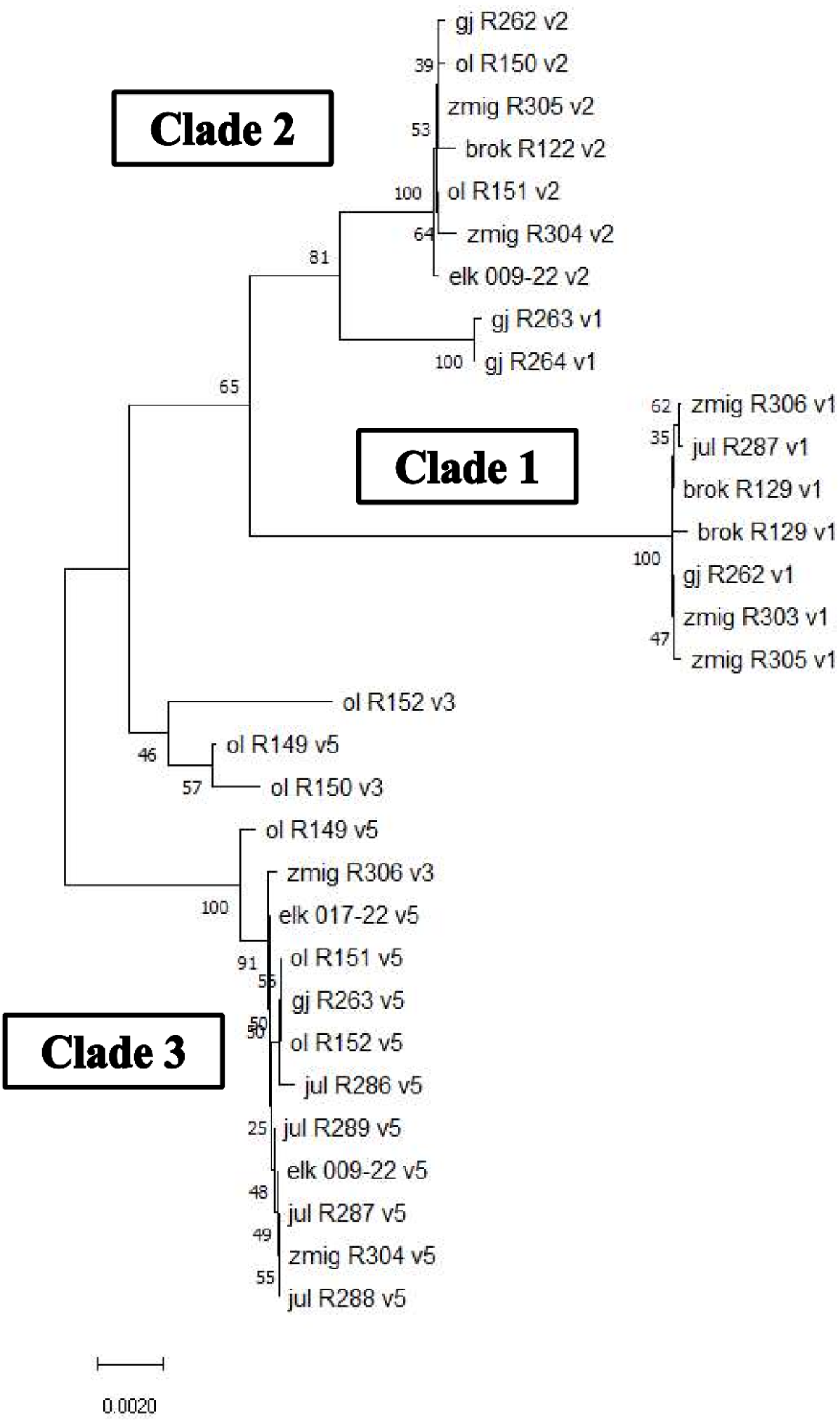
The full-length CFH sequences group into three main clades. Neighbour-joining tree of the full-length CFH sequences generated with MEGA11 using the Tamura 3-parameter model. The circles show the proportions of ancestry of the population of origin of the sequenced individual from the three genetic groups identified with RAD-seq SNPs (see Figure 4a). Sample IDs show population of origin, individual ID, and homology of the last exon with the CCP 20 variants (v1, v2, v3, or v5; see Figure 3A). gj = Goły Jon; ol = Olsztyn; zmig = Żmigród; elk = Ełk; jul = Julianka. Manual inspections of gj_R263_v1, gj_R264_v1, ol_R152_v3, ol_R149_v5, and ol_R150_v3 revealed that they are true recombinants between v2 and v1 (gj) or v2 and v3/v5 (ol).

**Figure 2:**
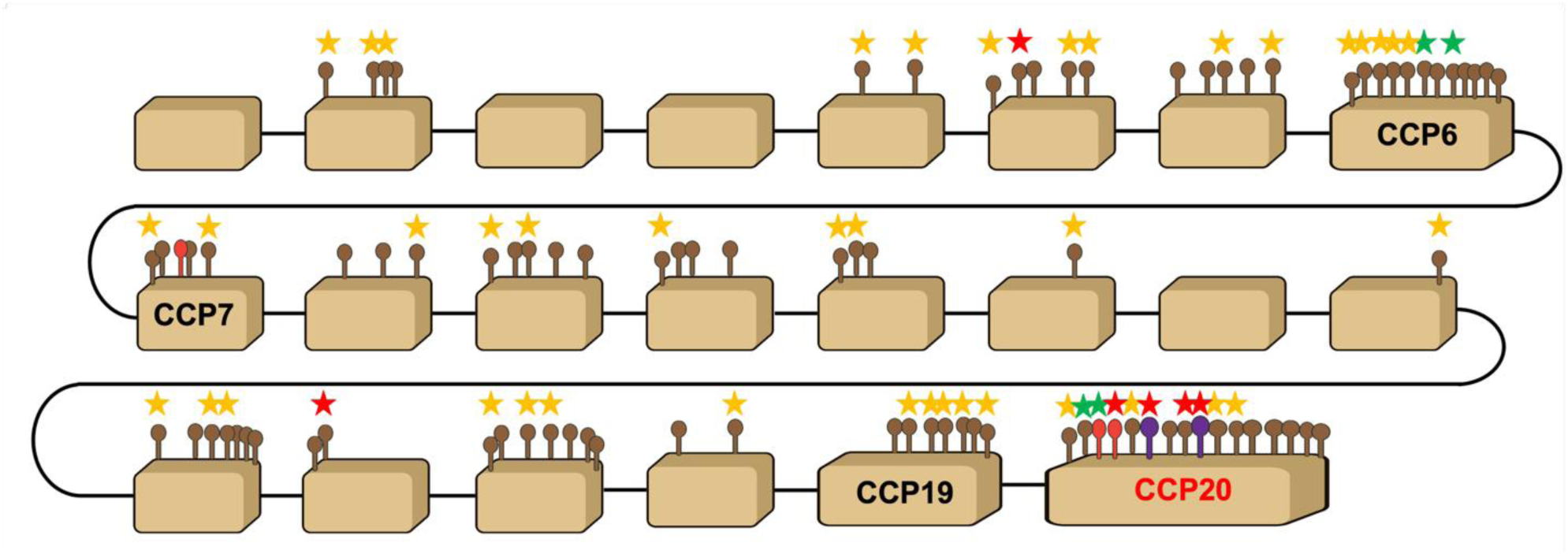
Many of the codons under positive selection (MEME and PAML) are in CCP 20 and overlap with or are adjacent to sites interacting with *Borrelia*. Schematic representation of the full CFH gene. Brown boxes represent each of the 22 exons; exons corresponding to CCP 6, 7, 19, and 20 are labelled. Lollipops above boxes represent non-synonymous SNPs and show their approximate location in the exon. Brown lollipops = non-synonymous SNP; red lollipops = non-synonymous SNP in a codon interacting with *Borrelia* or other pathogens (Cagliani et al., 2016; Meri et al., 2013); purple lollipops = non-synonymous SNP interacting with *Borrelia* and with the host’s polyanions (Meri et al., 2013). Yellow stars = codons under positive selection in MEME in either *C. glareolus* v1, v2, v5, or Node 5 (separating v2 from v1; Figure S2); red stars = codons under positive selection in PAML; green stars = codons under positive selection in both MEME and PAML. The positively selected sites associated to Node 4 (the branch leading to bank voles; Figure S2) are not shown.

We used PAML (Yang, 2007) to identify sites under pervasive positive selection across the phylogeny, and MEME to detect sites under episodic positive selection that may have occurred in the bank vole branches. For both analyses, we used one randomly selected CFH sequence from each of the three bank vole clades (v1, v2, and v5), four Cricetidae species, and murids *M. musculus* and *R. norvegicus* as an outgroup (Figure S2). The analysis with PAML revealed 16 sites with evidence of pervasive positive selection among rodents (Table 1). Within bank voles, MEME identified 46 sites with strong evidence of episodic positive selection (Empirical Bayes Factor >10; Table S1; Figure 2), 30 of which were also positively selected in other rodent branches (Dataset S1) while 16 were unique to bank voles. Most of the positively selected sites (27) were detected in the branch leading to the v1 CFH variant, while 12 and 5 sites were detected in the v2 and v5 branches, respectively. The remaining four sites were detected in the branch leading to the v1 and v2 clade (node 5). Fifty positively selected sites (different from the 46 above) were on the branch separating bank voles from other rodents (node 4). Both PAML and MEME identified six positively selected sites within CCP 20, although only two were in common between the two analyses (Table 1, S1). Furthermore, of the six sites identified by MEME within CCP 20, only one was significant in the branch leading to the bank vole clade, suggesting positive selection in this domain arose mostly after the bank vole lineages diverged. Many of the positively selected sites within CCP 20 in bank voles (identified with MEME, namely D1196, Y1207, I1213, and R1228) overlapped or were directly adjacent to sites involved in interactions with *Borrelia* (OspE and CspA), other pathogens, and/or self-components (Figure 2; Kraiczy et al., 2004; Meri et al., 2013; Cagliani et al., 2016). Interestingly, positively selected sites within CCP 20 involved in interactions with pathogens were found in all other rodent species, but they were different from those found in bank voles, except for Y1207, also found in *P. maniculatus* (Dataset S1). D1196 and Y1207 were also inferred by PAML, suggesting pervasive positive selection across the phylogeny.

**Table 1:** Sites under positive selection identified by PAML. Shading highlights codons in CCP 20.

| Site | Pr($\omega > 1$) | $\omega$ posterior<br>mean | SE |
| --- | --- | --- | --- |
| 219 | 0.962 | 2.735 | 0.66 |
| 223 | 0.95 | 2.709 | 0.684 |
| 352 | 0.999 | 2.817 | 0.572 |
| 362 | 0.991 | 2.8 | 0.594 |
| 367 | 0.978 | 2.773 | 0.627 |
| 390 | 0.97 | 2.756 | 0.644 |
| 405 | 0.995 | 2.809 | 0.584 |
| 575 | 0.951 | 2.711 | 0.681 |
| 948 | 0.999 | 2.818 | 0.571 |
| 1017 | 0.984 | 2.785 | 0.612 |
| 1047 | 0.973 | 2.762 | 0.638 |
| 1196 | 0.97 | 2.753 | 0.645 |
| 1207 | 0.982 | 2.782 | 0.615 |
| 1211 | 0.957 | 2.726 | 0.672 |
| 1217 | 0.999 | 2.818 | 0.572 |
| 1219 | 0.993 | 2.805 | 0.589 |
| 1221 | 0.998 | 2.815 | 0.575 |

Regarding other CFH domains, MEME detected 5 positively selected sites in CCP 19, 8 in CCP 6, and 2 in CCP 7 (Figure 2). Two sites in CCP 7 (I400 and E410) were significant in the branch leading to bank voles (Table S1) and were within a region bound by *Borrelia*’s CspA (Kraiczy et al., 2004).

Taken together, the results show that CFH in bank voles is highly polymorphic and subject to positive selection, particularly at the sites directly involved in interactions with pathogens.

### CCP 20 diversity shows a geographical pattern

We then focused on the CCP 20 domain, as this is the most polymorphic region and is functionally important for interactions with pathogens and self-molecules. We first amplified the fragment of 189 bp covering CCP 20 and then filtered co-amplifying CHF-related genes and possible pseudogenes by mapping sequences to the database of cDNA CCP 17-20 (*see above*). We found a total of six unique CCP 20 variants in our sample of 143 voles. All genotyped individuals had one or two CCP 20 CFH variants, confirming their allelic status. The phylogenetic analysis of the six CCP 20 alleles revealed three main clades corresponding to the main clades identified with the full sequence tree (Figure 1, 3a). The clades show a clear geographical pattern (Figure 3b). Clade 2 consists of three variants (v2, v4, and v6) and dominates in the SE, clade 1 contains only one variant (v1) and dominates in the NE, while clade 3, comprising two variants (v3 and v5), dominates in the SW. Populations from central regions show mixed frequencies of all clades.

**Figure 3:**
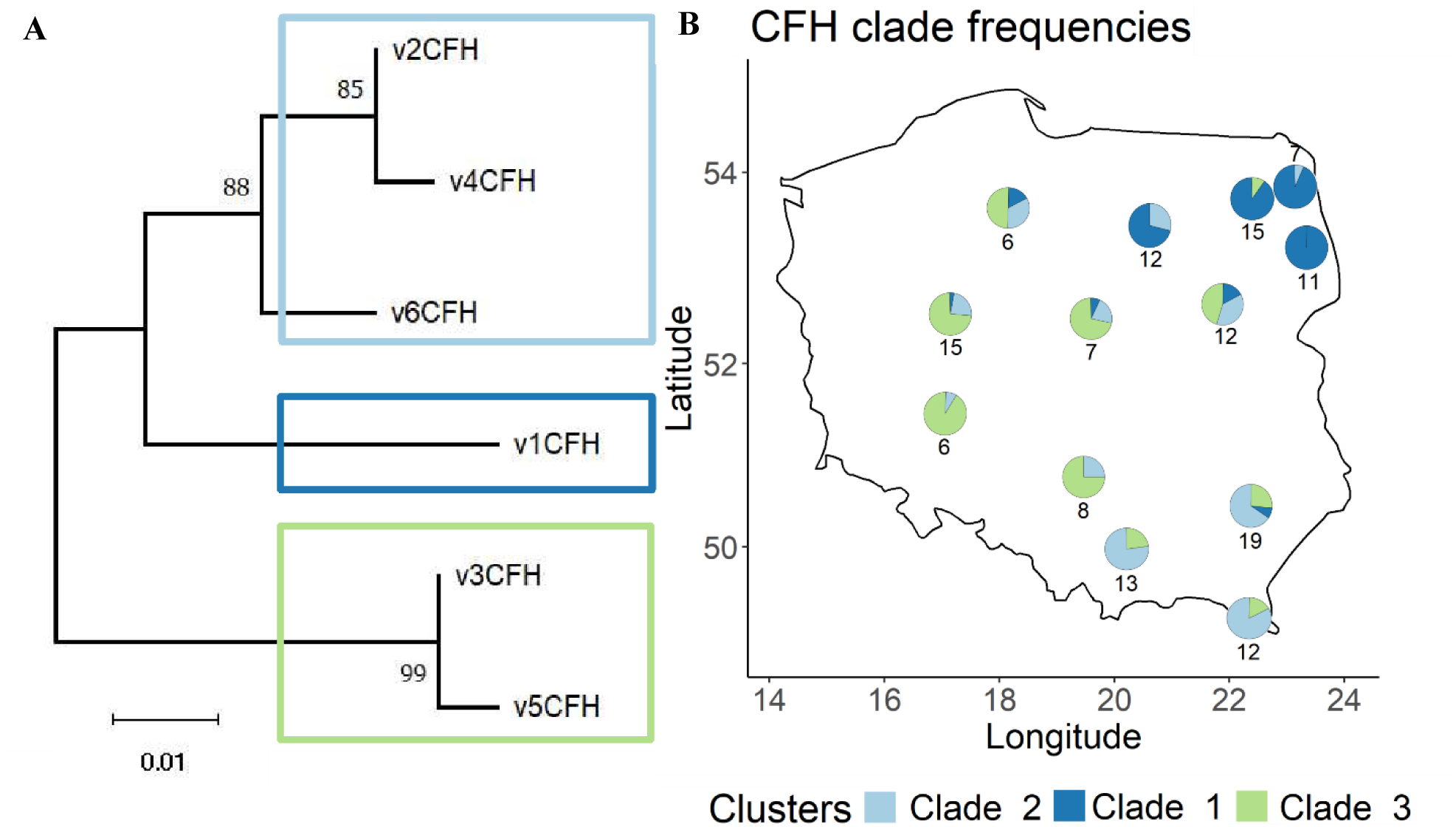
CFH is highly polymorphic with variants clustering in three clades with different frequencies across populations. (a) Neighbour-joining phylogenetic tree of the CFH variants at the CCP 20 domain. Numbers at nodes show bootstrap values. (b) Map of Poland with overlaid pie charts showing CFH clade frequencies per population. Numbers under (or above) the pie charts are sample sizes per population (number of sequenced individuals). Azure = clade 2 (v2, v4, and v6); blue = clade 1 (v1); green = clade 3 (v3 and v5). The same individuals as for RAD-seq were sequenced; however, CFH amplification failed for a few individuals.

### CFH shows a stronger population structure than the genomic background

Next, we compared the geographical pattern identified in CCP 20 with the overall bank vole population structure. To this aim, we conducted RAD sequencing on the same individuals screened for CCP 20 to obtain a reduced representation of the genome. SNP calling and genotyping of the RAD-seq data using Stacks resulted in 58,535 markers after filtering and thinning of closely positioned SNPs (average MAF after thinning = 0.219). Overall, all populations showed low differentiation (average pairwise F_ST_ = 0.032 ± 0.027 SE; Figure S3) and inbreeding (average inbreeding coefficient [F_IS_] = 0.017 ± 0.0023 SE; Table S2). Admixture analysis of population structure using RAD-seq markers revealed that populations cluster into three main groups (K = 3 had the lowest cross-validation error; Figure S4), and their structure is congruent with the geographical pattern observed in CCP 20 (Figures 3b, 4a). Central populations show larger degrees of admixture, consistent with their geographical locations (Figure 4a).

**Figure 4:**
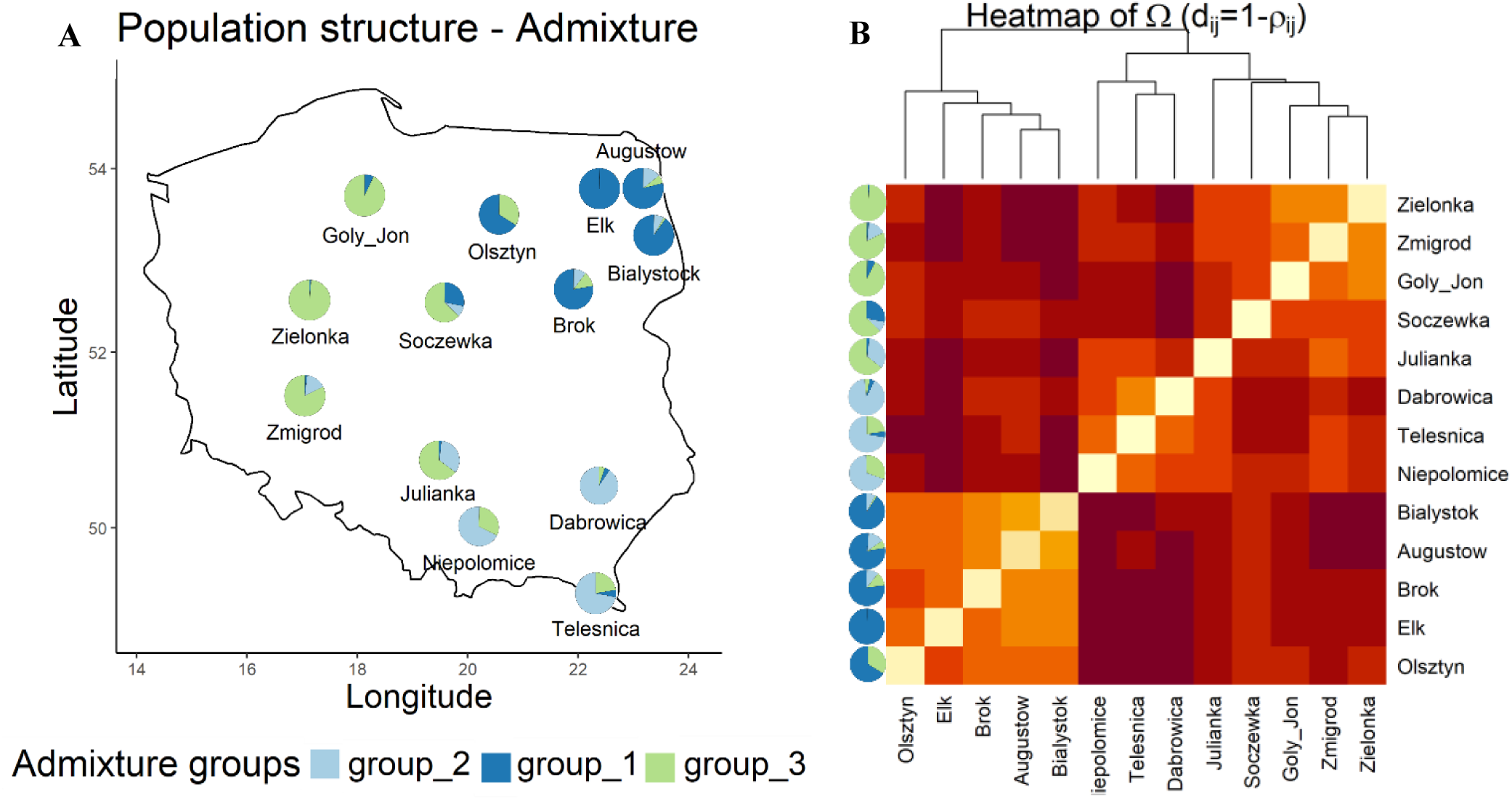
Bank vole populations in Poland cluster into three main groups. (a) Population structure identified with Admixture analysis of RAD-seq SNPs (K = 3). Pie charts show the admixture ancestry proportions (Q) averaged by each population. Azure: group 2 (∼Southern clade); blue: group 1 (∼Eastern clade); green: group 3 (∼Carpathian clade). Given that the 10 admixture runs with different seeds gave comparable results, Q values from one run chosen at random are shown. (b) Heatmap showing the population structure identified in the covariance (Ω) matrix. The lighter the colour of the cell, the stronger the correlation between the two populations. The Ω matrix was scaled to a correlation (similarity) matrix (ρ_ij_), and the dissimilarity matrix (d_ij_) between populations (i and j) was plotted. The tree was generated with hierarchical clustering using the average agglomeration method. The pies on the left are the same as in (a).

We then statistically assessed the congruence between genomic population structure and the geographical pattern of the CCP 20 variants, testing whether structure is stronger in CCP 20 compared to the genomic average. To do so, we conducted genomic scans of selection using the hierarchical, Bayesian method implemented in BayPass (Gautier, 2015). This is a robust outlier test that accounts for complex demographic histories, such as the one found in European bank vole populations that expanded from multiple glacial refugia (Wójcik et al. 2010, Marková et al. 2020). This method, therefore, does not make *a priori* assumptions on the demographic history of populations, which is instead incorporated in a covariance matrix (Ω) estimated from allele frequencies and reflecting the neutral correlation structure among populations. Genomic scans were conducted using the RAD-seq markers and the SNPs in CCP 20. Of the 20 SNPs in this domain, six were perfectly linked (they were adjacent and occurred in the same variants); therefore, they were clumped together in the analyses, which were therefore conducted on 16 variants.

The structure identified in the Ω matrix was congruent with that from Admixture (Figure 4b). The genomic scans revealed that frequencies of 13 of the 16 SNPs in the CCP 20 domain were outliers, showing stronger differentiation than expected from genome-wide data (Figure 5). Additionally, the RAD-seq dataset contained two SNPs located in intronic regions inside the CFH gene, of which one was found to be an outlier under diversifying selection. Therefore, CFH and, particularly, the CCP 20 domain are more structured than the genomic average.

**Figure 5:**
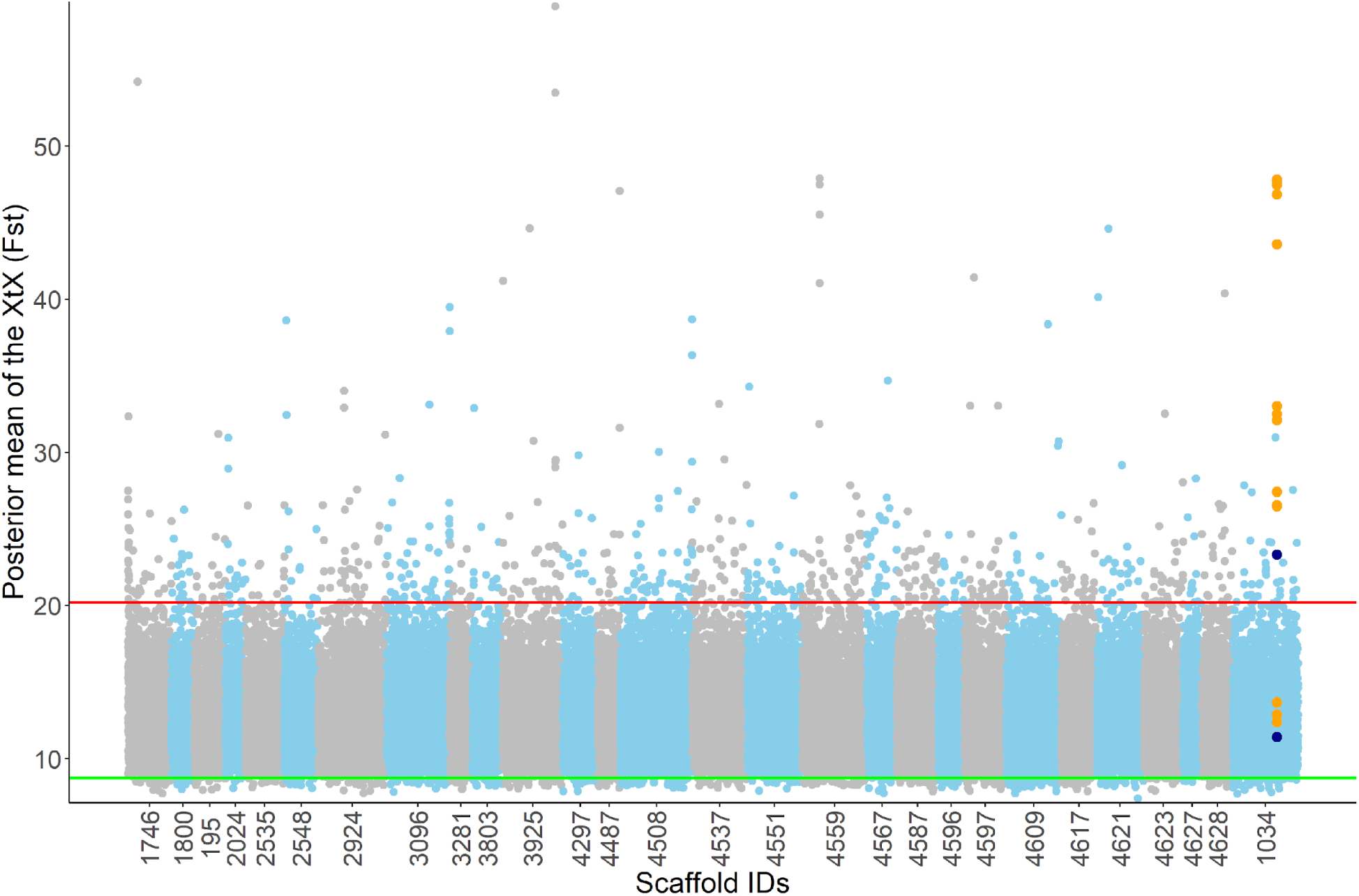
CFH SNPs are differentiated among populations and under diversifying selection. Manhattan plot showing the results of the genomic scans for signatures of selection. X-axis: cumulative genomic position in base pairs. The labels are the truncated scaffold IDs (the part after the ‘HRSCAF=’ in the labelling system for scaffolds used in the reference genome [Marková et al., 2023]); for graphical reasons, labels for the scaffolds with less than 2 SNPs are not shown, and their SNPs were combined with other scaffolds. Y-axis: mean from the posterior distribution of the X^T^X for each SNP. Each dot is a RAD-seq SNP; orange dots: SNPs in the CCP 20 domain; dark blue dots: the two RAD-seq SNPs inside the CFH gene. Scaffolds are shown with alternating colours. Red line: 99% quantile from the POD’s X^T^X null distribution; green line: 1% quantile from the same distribution.

### Outlier RAD markers are enriched for innate immunity and the alternative complement pathway

In addition to CFH, we identified 758 and 505 RAD SNPs showing extremely high or low differentiation, suggesting diversifying or balancing selection, respectively. We linked these to their closest genes and thus identified 272 and 196 unique, annotated genes under diversifying or balancing selection, respectively. The remaining outlier SNPs overlapped with, or were in proximity to, genes without annotations or located within retrotransposable elements.

Enrichment tests on the genes linked to SNPs under diversifying selection revealed 11 significantly enriched terms (Dataset S2; Figure S5a), including two immunity-related categories: the ‘complement activation, alternative pathway’ term (GO:0006957), to which CFH belongs, and ‘innate immune response’ (GO:0045087). The three outlier genes belonging to the first term were CFH, C3, and C5 (Dataset S2). Among the terms enriched for the SNPs under balancing selection, we found 12 significant terms (Dataset S3; Figure S5b), including ‘positive regulation of interleukin-6 production’ (GO:0032755) and ‘interleukin-6-mediated signalling pathway’ (GO:0070102).

Overall, the CCP 20 domain of CFH, interacting with pathogens and self-signals, is polymorphic and, together with other components of the alternative complement pathway, is under diversifying selection.

## Discussion

Complement factor H (CFH) is a key regulatory protein of the alternative complement pathway that is often hijacked by pathogens. Here, we describe a naturally occurring polymorphism in this gene and investigate molecular and population genetics signatures of selection acting on it in natural populations of the bank vole, one of the key European reservoirs of zoonotic pathogens, including *Borrelia afezelii* (Hanincová et al., 2003; Grzybek et al., 2020). We show extensive CFH polymorphism, particularly within domain 20 (CCP 20), involved in the protection of self-cells and serving as a binding site for pathogens, allowing immune evasion. Furthermore, this polymorphism is associated with signatures of natural selection in terms of (i) an excess of non-synonymous substitutions at several codons and (ii) elevated inter-population differentiation compared to the genome-wide average. Moreover, using RAD-seq, we identified additional outlier SNPs under diversifying selection in the proximity of innate immunity genes, including those involved in the alternative complement pathway.

### Polymorphism and selection on CFH

Polymorphism, coupled with signatures of positive selection, appears to be a common feature of genes involved in parasite recognition, including MHC genes essential for adaptive immunity (Hughes and Nei, 1988; Bernatchez and Landry, 2003) and Toll-like receptors (TLRs) of the innate immune system (Minias and Vinkler, 2022). This is thought to result from selective pressures exerted by parasites, whose molecular patterns evolve to evade recognition. Parasites are thought to adapt to the most common host genotypes; this can create selective pressures for amino acid substitutions in host receptor genes that restore binding and recognition, contributing to balancing selection and the maintenance of polymorphisms in immune genes (Bodmer, 1972; Ejsmond and Radwan, 2015; Migalska et al., 2022). This scenario is corroborated by more prevalent signals of balancing selection in immune receptor genes compared to downstream components of the immune response (Lundberg et al., 2020). However, whether the same mechanisms work within missing-self immune systems is not well understood. The regulators of missing-self immunity, such as CFH, are expected to be conserved because they recognise molecular markers of self (Carrillo-Bustamante et al., 2016); indeed, polymorphisms and mutations in CFH have been linked to autoimmune inflammatory disorders in humans (Ferluga et al., 2017).

Contrary to this expectation, here we found evidence of high polymorphism in CFH, confirming and extending results from Niedziałkowska et al. (2023), where two variants were identified in de novo transcriptomes from two populations. One explanation for this polymorphism is that CFH is under selective pressure from immune evasion by pathogens. Our analyses identified signals of positive selection along the branch leading to bank voles, including codons at sites in the CCP 20 domain adjacent to or overlapping with those involved in interactions with pathogen proteins, suggesting strong selective pressures to prevent pathogens from binding. Our results are remarkably similar to those reported for interspecific analyses of primate CFH (Cagliani et al., 2016), where signals of positive selection were detected in CCP 20 and in the *Borrelia* OspE codons interacting with CFH, lending further support to the host-parasite co-evolution hypothesis. The CCP 20 domain was under positive selection also in other rodents, but the sites targeted by selection differed among species. This suggests that selection in response to immune evasion is pervasive among mammals and that different sites within this domain may affect the binding of pathogen proteins.

To the best of our knowledge, a similar coevolutionary scenario was identified only in one other regulator of missing-self immunity before our study, the killer-cell receptor (KIR) that inhibits immune responses against cells presenting MHC class I molecules. This regulator effectively protects against viruses that suppress expression of MHC class I to prevent presentation of their antigen to T-cell receptors: the absence of MHC class I on these virus-infected cells makes them targets for clearance (Lanier, 1998; Parham, 2005). Despite targeting self-MHC, some representatives of the KIR family are polymorphic. A modelling study showed that this polymorphism can result from co-evolution with viruses producing decoy proteins that mimic MHC (Carrillo-Bustamante et al. 2015, 2016). Such coevolution would favour KIR variants that either escape targeting decoy proteins through increased affinity to their own MHC molecules or that become activators, rather than inhibitors, of immune responses. Similar to KIRs, CFH is a member of a large protein family, but the function of CFH-related proteins is incompletely understood (Pouw et al. 2015). However, several bacteria that have been identified to bind CFH are also able to bind to CFH-related proteins (Lambris et al., 2008; Kolodziejczyk et al., 2017), suggesting potentially complex coevolutionary scenarios that deserve further investigation.

The apparently adaptive polymorphism at CFH, particularly at CCP 20, is remarkable considering that this domain (along with CCP 7) is involved in binding to self-signals for immune recognition (Meri, 2016). This could possibly result in trade-offs between efficiency of binding to self-signals and avoidance of immune evasion. A recent theoretical study (Hummert et al. 2018) explored the role of such a trade-off in the maintenance of polymorphism at CCP 7, known in humans to be related to autoimmunity. The modelling showed that the trade-off may lead to the maintenance of stable polymorphism within populations if heterozygotes best balance infection resistance against autoimmunity. Whether the CCP 20 polymorphism leads to autoimmunity in bank voles deserves further investigation; however, we note that our results show higher population structuring at CFH compared to the genomic average, which is contrary to the predictions from heterozygote advantage, where the opposite pattern would be expected (Beaumont and Balding 2004).

The distribution of the CFH clades corresponded to the three genetic groups that we detected with RAD markers, suggesting both have been shaped by phylogeographic history. In addition to the two CFH divergent consensus haplotypes inferred from transcriptomes by Niedziałkowska et al. (2023), which dominate respectively in NE (clade 1) and SW (clade 3), we detected a third, previously undescribed CFH clade comprising the highest number of variants (clade 2) that dominated in SE. The three genomic groups, also identified by RAD-seq analysis in Niedziałkowska et al. (2023), although on a more limited geographic scale, likely reflect the history of postglacial colonisation (Kotlik et al., 2006; Wójcik et al., 2010; Marková et al., 2020). However, divergence of CFH major lineages is much higher than the genomic average. Indeed, the net synonymous divergence between our two most diverged bank vole CFH clades (clade 1 and 3) is 0.0184, which is about 4.6% of the average synonymous divergence between bank voles and yellow-necked mice immune genes (∼0.4; Zhong et al. 2021). Since the two species split 26.2 Mya (according to TimeTree.org; Kumar et al., 2022), we estimate that CFH accumulated divergence for about 1.2 My. This indicates that CFH polymorphism in bank voles predates the last glacial maximum (LGM) estimated to have occurred 18–23 kya (Mix et al., 2001) and suggests that the spatial patterning of CFH, consistent with genome-wide patterns, might have arisen via three non-exclusive scenarios. According to the first scenario, CFH could have diverged via selection throughout the Pleistocene, leading to local adaptation during glacial periods with no or little mixing during interglacial periods; however, it is not clear how to reconcile this scenario with the lower divergence of bank vole mitochondrial clades (30-70 kya; Wójcik et al., 2010), which is estimated to have occurred only during the last glaciation (ca. 115 kya). Alternatively, several CFH allelic lineages may have been maintained for a long time by balancing selection and then sorted differently in the glacial refugia. Extreme synonymous divergence of bank vole CFH clades compared to other expressed genes does indeed suggest long-term maintenance of the CFH lineages. While the higher CFH allele frequency differentiation compared to the genomic average does not support heterozygote advantage, balancing selection may have occurred via negative frequency-dependent selection (NFDS). NFDS may lead to rapid fluctuations in allele frequencies (e.g. Westerdahl et al., 2004; Migalska et al., 2022), and if host-parasite allele frequency cycles are out of phase in populations with limited gene flow, this can lead to higher population differentiation compared to neutral expectations, mimicking the signal of local adaptation (Spurgin and Richardson, 2010). Finally, in the third scenario, the geographic pattern of CFH would be due to admixture from an unknown source. Overall, our results indicate that selective forces, likely stemming from adaptation to pathogens, contribute to the inter-population differences in CFH allele frequencies. Whether selection on CFH is currently ongoing, possibly causing alleles to flow at a lower rate compared to the genomic average, deserves further investigation.

### Outliers at RAD-associated immunity genes

While our focus was on CFH, our analysis also identified several outliers in the RAD markers data (based on the X^T^X metric, an analogue of F_ST_ that accounts for the variance-covariance structure of the populations; Günther and Coop, 2013) enriched for various GO terms. Discussing all these outliers in detail is outside the scope of this study; here, we briefly discuss the outlying immunity genes. One of the terms that showed significant enrichment was ‘complement activation, alternative pathway’, which includes C3 and C5. C3 is a complex protein consisting of several subunits, each with different functions (Zarantonello et al., 2022; Geisbrecht et al., 2022). One subunit, C3b, binds to pathogen-associated molecular patterns (PAMPs) to induce opsonisation and further complement activation; the same subunit is also a target of immune evasion by *Staphylococcus aureus* in humans (Garcia et al., 2010; Geisbrecht et al., 2022). Additionally, mutations in C3 were linked to autoimmune disease (Zarantonello et al., 2022; Geisbrecht et al., 2022); therefore, population differentiation at this gene may result from a complex interplay of opposing selective pressures, i.e. increased binding efficiency to PAMPs, escaping binding from pathogens, and evolutionary constraints due to autoimmune disease. C5 is a downstream component of complement that is also the target of immune evasion from pathogens from different taxa (bacteria, ticks, fungi; Laursen et al., 2012; Luo et al., 2013). Both CFH and C5 interact with C3, and the specificity of their interaction may possibly contribute to population differentiation. Enrichment of complement cascade genes among those subject to local adaptation contrasts with findings from Nandakumar et al. (2023). This study reports significant signatures of balancing selection acting on four of the 44 complement cascade genes investigated, based on β population statistics reflecting a build-up of alleles at similar frequencies. These four genes did not include those we found to be subject to diversifying selection, so our data highlight that different components of the complement system seem to be involved in different co-evolutionary dynamics. This is not unexpected given that host-parasite co-evolution can take the shape of both balancing selection and recurrent selective sweeps (Woolhouse et al., 2002).

Additional immune genes closest to the SNPs with evidence for diversifying selection in our study include CD180 antigen, Annexin A1 (ANXA1), Ankyrin repeat domain 17 (ANKRD17), Interferon-induced protein with tetratricopeptide repeats 1 (IFIT1), and Interleukin 1 receptor accessory protein (IL1RAP). CD180 (a Toll-like receptor) and IFIT1 bind to PAMPs (McDougal et al., 2024; Xie et al., 2025), with IFIT1 also being a target for immune evasion (Xie et al., 2025). Thus, because of their function, these genes represent potential candidates for host-pathogen coevolution. Polymorphisms in IL1RAP, ANKRD17, and ANXA1 are associated with disease (Yu et al., 2020; Vilkeviciute et al., 2020; Tinatin et al., 2023; Granell et al., 2023). These genes are involved in immunity as signal transducers, interacting with hosts’ receptors, with no evidence so far that they could be targets of immune evasion. Immunity genes were also represented among the outlier loci under balancing selection, including enrichment of loci involved in the Interleukin 6 signalling pathways. Interestingly, among the genes in this pathway, we found TLR6, which was the only TLR without evidence of balancing selection (based on Fu and Li’s D* and Tajima’s D) in the bank vole populations investigated in Kloch et al. (2018). The reasons for the differences between these findings require further investigation.

## Conclusions

Our study revealed extensive polymorphism in CFH, particularly in CCP 20, which included several candidate sites under positive selection. Coevolution with pathogens whose surface proteins bind to this domain to evade destruction by the complement cascade seems the most likely explanation for this result, further supported by population genetic evidence for diversifying selection acting on CCP 20. Importantly, our findings provide novel evidence that genes involved in missing-self immunity, such as CFH, which are typically thought to be evolutionarily constrained due to their essential roles in self–recognition, can also be targets of adaptive evolution. The consequences of this naturally selected polymorphism on host fitness need to be addressed in future studies. More broadly, this work indicates that genes involved in innate immunity, particularly in the alternative complement pathway, are among the most differentiated across populations, likely reflecting selective pressures imposed by pathogens.

## Material and Methods

### Sampling

To capture bank voles’ representative diversity, sampling was conducted at 13 locations across Poland, spanning the contact zone between the two major phylogeographic lineages, Carpathian and Eastern, as identified by mitochondrial DNA (Wójcik et al., 2010; Marková et al., 2020; Table S3; Figure S6). We collected ear biopsies from *C. glareolus* in 2020-2023 during August-September under approval of the Local Ethics Committee for Animal Experimentation in Poznań, decision no. 35/2021. The samples were promptly stored in ethanol and frozen until DNA extraction. A subsample was preserved in RNAlater Stabilisation Solution (Thermo Fisher Scientific, Waltham, MA, USA) for RNA extraction. Samples from Teleśnica and Niepołomice were provided by Prof. Paweł Koteja (Jagiellonian University, Kraków, Poland) and were collected in 2009 (under approval of the local Ethics Committee in Kraków, Decision No. 48/2007). DNA was extracted from bank vole ear samples with the NucleoMag 96 Tissue Kit (Macherey-Nagel, Duren, Germany) following the manufacturer’s protocol. Total RNA was extracted with the ReliaPrep RNA Tissue Miniprep System (Promega, Madison, WI, USA). cDNA was synthesised using Maxima H Minus First Strand cDNA synthesis Kit (Thermo Fisher Scientific).

### CFH genotyping-by-sequencing

To obtain full-length CFH sequences, we used primers CCP0-F: 5’-ATGAGACTATCAGCAAGAATTATCTGGCT-3’ and CCP20-R: 5’-CATGAATATTTCTCAACAAAKRTTCA-3’, amplifying cDNA of a total length of 3,780 bp spanning exons 1 to 22. These primers were designed based on *de novo* assembled transcripts from Niedziałkowska et al. (2023) and Różańska-Wróbel et al. (2025) aligned with *M. musculus* transcripts (NCBI Reference Sequence: NM_009888.3) for exon identification. We used these primers to amplify CFH sequences from the cDNA of 19 individuals from 9 populations across Poland (Table S4), which were then sequenced using ONT MinION with the V14 rapid barcoding kit. Consensus sequences were obtained with the SLANG pipeline (Dorfner et al., 2022), which uses VSEARCH (Rognes et al., 2016) to cluster reads within and then between samples based on similarity thresholds. Since we wanted to identify different alleles from a single locus, we used very stringent similarity thresholds, i.e., 0.97 and 0.99 for within- and between-sample clustering, respectively. All other settings were set to default. We selected and manually inspected the CFH consensus sequences with a length >3,400 bp and supported by the largest number of reads.

We then focused on the most polymorphic CCP 20 domain of CFH. Based on full-length sequences, we designed primers 20F and 20R (CCP20-F: 5’-CATGTGTAATATCAGAAGAGAYCA-3’ and CCP20-R: 5’-CATGAATATTTCTMAACAAAGGTTCAT-3’) to amplify CCP 20 from genomic DNA. The primers were used to amplify CFH CCP 20 from the same samples as for RAD-seq; however, 11 amplifications were unsuccessful. The primers were barcoded to identify unique samples as described by Kozich et al. (2013), except that the ’N’ (0-3 nucleotide) spacers were added to increase library diversity during Illumina sequencing. PCR was performed using the Type-it Microsatellite PCR Kit according to the manufacturer’s protocol. The reaction setup contained 1 µL of template DNA (∼100 ng), 1 µL of CCP20-F primer (final concentration: 0.4 µM), 1 µL of CCP20-R primer (final concentration: 0.4 µM), 12.5 µL of 2x Type-it Multiplex PCR Master Mix, and nuclease-free water up to 25 µL. Thermal cycling was performed with an initial denaturation at 95 °C for 5 minutes, followed by 28 cycles of denaturation at 95 °C for 30 seconds, annealing at 58 °C for 90 seconds, and extension at 72 °C for 30 seconds, with a final extension at 60 °C for 30 minutes. CFH amplicons were sequenced using the Illumina MiSeq platform with the MiSeq Reagent Micro Kit v2 (paired-end, 2 × 150 bp). The resulting reads were merged using AmpliMERGE and analysed with AmpliSAS for demultiplexing, clustering, and filtering of unique variants (Sebastian et al., 2016). We used default settings for Illumina reads and enabled the ’discard frameshifts’ option.

We obtained up to 11 variants per individual (instead of the expected two), suggesting that the CCP 20 primers, in addition to CFH, also amplified CFH-related genes and possibly pseudogenes. To exclude these non-CFH genes, we first created a database of expressed CFH, obtained by sequencing cDNA from a representative sample of 50 individuals from 9 populations (Table S5). Because CFH-related genes do not have CCP 17, and most of them also miss CCP 18 (Pouw et al., 2015), we amplified cDNA in a nested PCR using forward primers CCP17-F (5’-TGATTGTGACAGTTTACCCAAGTATGA-3’, 10 cycles) and CCP18-F (5’-TGATGTGTCAAAATGGGATTTG-3’, 20 cycles) and the reverse primer CCP20-R to selectively exclude CFH-related genes. The amplicons were sequenced using the Illumina MiSeq platform with the Reagent Kit v3 (paired-end, 2 × 300 bp). Since we expected to obtain sequences from a single expressed locus, we checked and confirmed that each individual possessed only one or two alleles. To exclude CFH-related genes and pseudogenes, we then mapped the CCP 20 sequences obtained from genomic DNA to the database of seven expressed CFH sequences; matching CCP 20 sequences (six variants) were retained for downstream analysis. All individuals in our sample had one or two variants of the filtered sequences, confirming their allelic status.

Additionally, we reanalysed transcriptomic data originating from western (Włocławek) and eastern (Białowieża) Poland (Niedziałkowska et al., 2023). Raw reads were filtered using Trimmomatic (Bolger et al., 2014) and mapped to the reference transcriptome with BWA-MEM (Li, 2013). Duplicate reads were marked with samtools markdup, and reads with a mapping quality below 20 were removed. Genotypes were called across all sites using bcftools, and positions with a quality score or a mean mapping quality below 30 were excluded. For downstream analyses, we retained only genotypes with a minimum depth of 6. Using a custom Python script, we calculated synonymous and nonsynonymous divergence between the eastern and western populations following the method of Nei and Gojobori (1986). We also estimated within-population nucleotide diversity at synonymous and nonsynonymous sites, as well as net divergence. All analyses were performed per protein-coding gene, and the resulting values for the predicted CFH gene were compared against the genome-wide distribution obtained from all genes.

### Positive selection on CFH (d_N_/d_S_ analyses)

To identify signatures of positive selection, we conducted nonsynonymous-to-synonymous substitution (d_N_/d_S_) analyses on the full CFH transcripts. To obtain CFH sequences from related species, we used blastn to search for CFH in published Iso-seq databases from Muridae and Cricetidae (*Cricetulus griseus*: DRR568657-DRR568660; *Mus spicilegus*: ERR9929550-ERR9929561; *Mus spretus*: ERR9929562-ERR9929573; *Mesocricetus auratus*: SRR12589345, SRR14718534, SRR18449643; *Peromyscus maniculatus*: SRR29268902; *Microtus pennsylvanicus*: SRR30640723, SRR30640724). As a query, we used one of our full CFH sequences for all databases (‘jul_R289_v5’) and the mouse CFH (‘BC066092-1’) for Muridae databases. An e-value threshold was set at 1e^-30^. We then filtered the results to retain identified sequences with an alignment length greater than 3,000 bp; for each species and accession, we selected the sequence with the highest bitscore. Finally, sequences were extracted from fastq files with seqkit grep (Shen et al., 2016). Manual checks revealed a high degree of divergence between accessions within both *M. spicilegus* and *M. spretus* that was comparable to divergence between species, suggesting extensive sequencing errors. This result was obtained both when using sequences blasted from bank vole or mouse CFH. We therefore discarded those species from the following analyses. For the species with multiple accessions, the sequence with the highest bitscore and length was chosen. CFH CDSs for *M. musculus* (‘BC066092-1’) and *R. norvegicus* (‘NM_130409-2’) were available and therefore directly downloaded. The d_N_/d_S_ analyses were therefore conducted on Cricetidae, *M. musculus*, *R. norvegicus*, and three bank vole variants, one from each of the three clades (‘jul_R289_v5’, ‘gj_R262_v1’, ‘brok_R122_v2’). These variants were added because we estimated they are old (1.2 My, *see Results*) and likely evolved independently for long periods of time in glacial refugia, where they reached high frequencies. Results of d_N_/d_S_ analyses using only one variant were comparable.

Sequences were aligned with Macse (Ranwez et al., 2011), trimmed to the same length, and deprived of the stop codon. This alignment was used to generate a phylogenetic tree with MEGA11 (Tamura et al., 2021) using the neighbour-joining method tested with bootstraps (1000 replications) and the Tamura 3-parameter DNA evolution model (Figure S2). We then conducted tests of d_N_/d_S_ with PAML (Yang, 2007) and MEME (Murrell et al., 2012). We ran PAML under all site models (M0, M1a, M1b, M7, and M8) using a mutation-selection model with observed codon frequencies as estimates. Significant differences between nested models were checked using likelihood ratio tests, i.e., M0 vs M1a, M1a vs M2a, and M7 vs M8. We considered positively selected sites those with a probability of ω >1 greater than 0.95.

MEME was run with default settings. We considered as sites with evidence of positive selection those with an Empirical Bayes Factor >10 in the three bank vole tree leaves, in ‘node 4’ (leading to the bank vole clade), and in ‘node 5’ (leading to bank vole v1 and v2).

The average sequence divergence between all full-length CFH sequences was calculated with the *nuc.div()* function from ‘pegas’ (Paradis, 2010), which uses Nei’s (1987) formula.

### RAD sequencing protocol

To obtain a genomic background for variation and genetic structuring, the DNA from all samples genotyped for CCP 20 was sequenced with the 3RAD approach (Bayona-Vasquez et al., 2019), a reduced-representation library sequencing method. It is based on cutting DNA with a set of restriction enzymes, resulting in a common fraction of loci across samples, which allows for the identification of single-nucleotide polymorphism (SNP) markers. Libraries were constructed following the protocol provided in Bayona-Vasquez et al. (2019), with the following differences: i) Illumina indexed primers were incorporated in a single PCR, with 12 cycles; ii) purification at all steps was performed using AMPure XP magnetic beads (Life Sciences), following manufacturer’s protocol; iii) selected size range was 300-600 bp. The size range of the DNA fragments for RAD sequencing was chosen based on the results of the *in-silico* digestion of the *C. glareolus* genome (accession: GCF_902806735.1) using the three enzymes of the 3RAD protocol (*XbaI*, *EcoRI*, and *NheI*) and a modified Python script from Driller et al. (2020). The results showed that the 300-600 bp size range would yield 20,000-30,000 loci for population genomic inferences (Figure S7).

Samples were divided into three pools, and a set of adapters including internal indexes (i5 and i7 adapters from Table S3 in Bayona-Vasquez et al. [2019]) was individually ligated to samples in each pool. Each one and a half pool received a different combination of Illumina indexes (iTru5_001_A-iTru7_101_01 and iTru5_001_B-iTru7_101_02 from Table S3 from Bayona-Vasquez et al. [2019]). Combining the half pools with the full pools containing the alternative combination of Illumina indexes allowed us to perform the paired-end sequencing (150 cycles) of all samples in two lanes of NovaSeq 6000 SP flow cell (Illumina). An addition of 20% PhiX was used to counterbalance the effect of fixed enzyme restriction sites.

### SNP calling and genotyping

RAD-seq data processing and loci building and genotyping were conducted with the Stacks pipeline (Catchen et al., 2013; Rochette and Catchen, 2017) and other tools operating in R v.4.2.2 (R Core Team, 2022). Firstly, we used the process_radtags program in Stacks to demultiplex the raw reads from the Illumina sequencing run (barcodes in line with sequence on both paired-end reads), remove any read with an uncalled base, discard reads with average quality scores (phred) within any sliding window <10 (window size = 15% of the length of the read), truncate reads to a length of 140 bp, and filter the Illumina adapters with 1 mismatch allowed in the adapter sequence. We assessed the quality of each sequenced sample and generated summary reports using the fastqcr package (Andrews, 2010; Kassambara, 2023). Samples with notably low coverage, containing fewer reads than 10% of the median of all samples (302,396.5 reads), were dropped (65 samples; Figure S8).

Following Paris et al. (2017), before running the Stacks pipeline, we conducted optimisations of the three Stacks parameters that define initial loci and visualised the results using ‘RADStackshelpR’ (DeRaad, 2021a). ‘vcfR’ (Knaus and Grunwald, 2017) was used to calculate summary statistics from output vcf files. First, *m* (the minimum coverage to create a stack) was iterated across 3-7, with default *M* and *n*; then, *M* (the maximum distance allowed between stacks within a locus) was iterated across 1-8, with optimal *m* (Figure S9) and default *n*; finally, *n* (the number of mismatches allowed between samples to build catalogs) was iterated across 2-4 with optimal *m* and *M*. Optimizations were conducted on 16 samples chosen at random and keeping all other parameters as the final Stacks run (*see below*). This process revealed that *m* = 3, *M* = 3, and *n* = 4 were optimal values for the structure of our data (Figure S9).

The RAD-seq reads were assembled using the *de novo* Stacks pipeline. Because the *in-silico* digestion showed the presence of paralogous loci, we implemented a strict haplotyping approach to prune those loci, following Piertney et al. (2023). The ustacks program was run with *m* = 3, *M* = 3, two maximum stacks per single *de novo* locus, nine maximum mismatches allowed to align secondary reads to primary stacks, and a bounded model with an upper bound for the error rate (epsilon) of 0.05. The cstacks, sstacks, tsv2bam, and gstacks programs were run with default parameters, except that *n* = 4 was used in cstacks and gapped alignments between stacks were disabled in both cstacks and sstacks. After these steps, we removed from subsequent analyses 43 samples that showed low coverage (<10x; Figure S10a) and one outlier sample with a low number of genotypes (Figure S10b). We then integrated the *de novo* assembled loci with their alignment positions to the genome. Each locus was aligned to the reference genome (Marková et al., 2023) using BWA mem (Li, 2013) and the output was converted to BAM using samtools (Li et al., 2009). The stacks-integrate-alignments script was used to inject the alignment coordinates back into the Stacks output.

A final check was conducted at this stage to remove remaining paralogous loci. We first ran the populations program in Stacks to generate a filtered VCF file with the loci present in at least 50% of the individuals for each population, a Minor Allele Frequency (MAF) of 0.01, and a maximum observed heterozygosity of 0.6. ‘HDPlot’ (McKinney et al., 2017) was then run on the vcf file to identify the SNPs with an observed read ratio deviating from the expected for heterozygotes (1:1). We considered as putative paralogs all loci with at least one SNP with an absolute observed read ratio (|D|) above a stringent threshold of 6 (Figure S11). Additionally, we used VSEARCH to cluster loci by sequence similarity; all loci clustering with other loci were regarded as putative paralogs (as in Piertney et al., 2023). VSEARCH was run with an identity threshold of 0.1 (obtained via optimization across 0.1-0.9; the 0.1 identity threshold generated the least number of clusters, providing a conservative set of singleton loci; Piertney et al., 2023) with identity defined as the (matching columns) / (alignment length), no masking regions, rejection of the sequence match if the alignment begins with or contains gaps, and a complete database search. The paralogous loci identified by both methods (3,600; Figure S11) were blacklisted and removed from further analyses.

The population program was run again with the same options as before, except the paralogous loci were excluded from the report, and the MAF was 0.03. Finally, we conducted quality and completeness filtering of the SNP dataset using ‘SNPfiltR’ (DeRaad, 2021b) and ‘vcfR’. First, we removed genotypes supported by a read depth below five, quality below 30, and a heterozygote allele balance outside of the 0.1-

0.9 range. We then applied a maximum depth cutoff to remove all SNPs with depth higher than double the mean depth of all SNPs (maximum depth cutoff = mean depth [60x] * 2 = 120x). Two samples had missing data >50% and were removed from further analyses. We then applied an 80% SNP completeness cutoff (i.e., 20% of missing data allowed per SNP) as a trade-off between minimising the overall proportion of missing data across samples and maximising the total number of loci (Figure S12). Duplicated SNPs (SNPs that were present twice in the VCF file) were also removed by retaining the one with the lowest amount of missing data. To reduce linkage disequilibrium between markers, the final VCF file was thinned to one SNP with the highest MAF for each RAD locus using custom scripts and VCFtools (Danecek et al., 2011). Finally, since ’Scaffold_1610;HRSCAF=1701’ blasted to the mouse X chromosome, all SNPs in this scaffold (2,513) were removed to avoid ploidy differences between male and female individuals that may inflate subsequent population genomic analyses.

A total of 58,535 RAD SNP loci passed filtering and thinning, for a total of 154 remaining samples.

The number of remaining samples for each of the 13 populations is presented in Table S4.

### Population structure

Genetic population structure was analysed with the Admixture program (Alexander et al., 2009; Alexander and Lange, 2011) using SNPs from RAD-seq. The filtered and thinned VCF file was converted to bed, and the additional files required, using PLINK (Chang et al., 2015). Admixture was run 10 times with random seeds and K = 1-13 to find the value with the lowest cross-validation error. For each value of K and each run, results were summarised and visualised using Pong (Behr et al., 2016).

### Genome-wide scan for selection and enrichment tests

We conducted genome-wide scans for adaptive genetic differentiation across populations using the Bayesian hierarchical method implemented in the BayPass program (Gautier, 2015). BayPass was run with default settings on the RAD and CFH (CCP 20) SNP allele counts dataset. Genotype information was extracted for each RAD SNP and sample from the filtered and thinned VCF file using VCFtools and formatted for BayPass with the *gt.format* function in ‘rCNV’ (Karunarathne et al., 2023). This was complemented with the CFH allele counts calculated for the biallelic SNPs identified in the CCP 20 domain (*see Results*). Six of the 20 SNPs in this domain were adjacent and occurred in the same variants and were therefore clumped together, for a total of 16 SNPs used in the analyses. To carry out our focal test on SNPs within CCP 20, we included all biallelic SNPs in this domain, rather than thinning them as we did for RAD markers. This approach is more robust compared to focusing on just the one SNP within CCP 20 that would remain after thinning; nonetheless, the one SNP within CCP 20 with the highest MAF (MAF = 0.5), which we would use in the analysis if we conducted thinning of CCP 20 SNPs with the same rationale as for RAD markers, was an outlier.

We first estimated the Ω matrix and the X^T^X statistic (an F_ST_-like statistic that accounts for shared population history; Günther and Coop, 2013). Then, to identify X^T^X outliers, we simulated a pseudo-observed dataset (POD) containing 100,000 SNPs using parameters fixed to the posterior means of the original data. The POD was then analysed with BayPass using the same settings as before to generate a null distribution of the X^T^X values. This distribution was then used to define 99% and 1% quantile thresholds. We considered as outliers under diversifying selection all SNPs with X^T^X values above the 99% quantile of the null (POD) distribution, and SNPs under balancing selection, those with X^T^X values below the 1% quantile.

To perform functional enrichment on the outlier SNPs (functional annotation clustering), we first used bedtools closest (Quinlan and Hall, 2010) to search for the genes in the reference genome overlapping or nearest to each outlier SNP under diversifying or balancing selection; we then extracted the corresponding UniProt gene IDs from the annotations file and mapped them to the UniProtKB database on the UniProt website (https://www.uniprot.org/id-mapping). This list of gene IDs was used to conduct enrichment tests with David (Sherman et al., 2022). We used as background for the analysis the nearest or overlapping genes to all RAD SNPs (5,951 genes), which were extracted as explained above.

## Author contributions

RFN and JR designed the study and wrote the manuscript; all other authors contributed to the development and revision of the manuscript. RFN conducted all bioinformatic analyses; MHR and MK provided bioinformatic consultation and revised the manuscript; MHR conducted 3RAD library preparation, and JRW and KP performed the remaining molecular work. PK provided genomic resources. JR, JRW, RFN, KP, and MK collected samples.

## Acknowledgments

We thank Prof. Paweł Koteja for providing samples from Teleśnica and Niepołomice; Dr Larissa Arantes for providing modified scripts to conduct *in-silico* digestion with three enzymes; Dr Agnieszka Szubert-Kruszyńska for administrative organisation; Józefina Wasilewska and Dr Aleksandra Łukasiewicz for their help during sampling; and Dr Julia Zaborowska for conducting transcriptomic analyses of bank voles that we used to design CFH primers. This research was supported by the National Science Centre, Poland (Grant/Award Number: 2019/34/A/NZ8/00231). PK was supported by MEYS CZ (project LUAUS25009).

## Data availability statement

All data will be made available in FigShare upon publication of the manuscript.

