## Supplementary Tables and Figures for "A key regulator of missing-self innate immunity is polymorphic and under diversifying selection"

**Table S1:** Sites under episodic positive selection identified by MEME. Results were subset to include only bank vole leaves (*C. glareolus* v1, v2, and v5), the node leading to bank voles (Node 4), and the internal node separating v1 and v2 from v5 (Node 5). EBF = Empirical Bayes Factor. Shading highlights codons in CCP 20.

| Site | Branch | EBF | Site | Branch | EBF | Site | Branch | EBF |
| --- | --- | --- | --- | --- | --- | --- | --- | --- |
| 41 | M_glareolus_v1 | 177.61 | 648 | M_glareolus_v2 | 42.31 | 400 | Node4 | 30.87 |
| 68 | M_glareolus_v1 | 330.47 | 1040 | M_glareolus_v2 | 325.34 | 410 | Node4 | 182.18 |
| 70 | M_glareolus_v1 | 257.37 | 1166 | M_glareolus_v2 | 1129.72 | 418 | Node4 | 166.68 |
| 152 | M_glareolus_v1 | 12.44 | 1213 | M_glareolus_v2 | 2570.77 | 450 | Node4 | 481.97 |
| 209 | M_glareolus_v1 | 18.31 | 1233 | M_glareolus_v2 | 71.72 | 528 | Node4 | 160.1 |
| 239 | M_glareolus_v1 | 339.41 | 187 | M_glareolus_v5 | 6030.05 | 531 | Node4 | 63.37 |
| 243 | M_glareolus_v1 | 196.33 | 291 | M_glareolus_v5 | 20.99 | 570 | Node4 | 24.86 |
| 341 | M_glareolus_v1 | 169.05 | 342 | M_glareolus_v5 | 22.15 | 601 | Node4 | 36.47 |
| 345 | M_glareolus_v1 | 7936.03 | 414 | M_glareolus_v5 | 27.56 | 689 | Node4 | 70.25 |
| 380 | M_glareolus_v1 | 1983.44 | 1144 | M_glareolus_v5 | 5099.12 | 714 | Node4 | 4130.34 |
| 392 | M_glareolus_v1 | 122.45 | 17 | Node4 | 30.22 | 716 | Node4 | 235.77 |
| 497 | M_glareolus_v1 | 744.93 | 218 | Node4 | 140.2 | 717 | Node4 | 165 |
| 521 | M_glareolus_v1 | 21.48 | 225 | Node4 | 42.01 | 737 | Node4 | 378.71 |
| 573 | M_glareolus_v1 | 162.24 | 263 | Node4 | 62.33 | 747 | Node4 | 43.19 |
| 737 | M_glareolus_v1 | 117.94 | 269 | Node4 | 6868.85 | 802 | Node4 | 54.14 |
| 863 | M_glareolus_v1 | 55.15 | 271 | Node4 | 53.02 | 807 | Node4 | 3581.35 |
| 872 | M_glareolus_v1 | 25.83 | 278 | Node4 | 38.91 | 823 | Node4 | 50.51 |
| 924 | M_glareolus_v1 | 982.1 | 280 | Node4 | 341.72 | 837 | Node4 | 60.91 |
| 931 | M_glareolus_v1 | 3108.21 | 289 | Node4 | 156.25 | 951 | Node4 | 27.38 |
| 1006 | M_glareolus_v1 | 4768.46 | 295 | Node4 | 40.43 | 956 | Node4 | 180.54 |
| 1026 | M_glareolus_v1 | 25.96 | 299 | Node4 | 16.88 | 960 | Node4 | 61.76 |
| 1091 | M_glareolus_v1 | 768.89 | 301 | Node4 | 896.84 | 986 | Node4 | 61.97 |
| 1156 | M_glareolus_v1 | 49.69 | 308 | Node4 | 62.95 | 1047 | Node4 | 40.27 |
| 1161 | M_glareolus_v1 | 1217.66 | 311 | Node4 | 24.68 | 1049 | Node4 | 64.45 |
| 1196 | M_glareolus_v1 | 39.64 | 323 | Node4 | 29.04 | 1069 | Node4 | 2196.67 |
| 1207 | M_glareolus_v1 | 227.1 | 338 | Node4 | 333.11 | 1082 | Node4 | 33.31 |
| 1228 | M_glareolus_v1 | 44.26 | 362 | Node4 | 16.64 | 1117 | Node4 | 28.95 |
| 317 | M_glareolus_v2 | 88.4 | 369 | Node4 | 97.63 | 1138 | Node4 | 216.33 |
| 336 | M_glareolus_v2 | 140.05 | 383 | Node4 | 979.53 | 1151 | Node4 | 95.84 |
| 347 | M_glareolus_v2 | 11.16 | 384 | Node4 | 92.57 | 1233 | Node4 | 271.84 |
| 352 | M_glareolus_v2 | 11.36 | 390 | Node4 | 12.89 | 1156 | Node5 | 11.38 |
| 362 | M_glareolus_v2 | 62.94 | 391 | Node4 | 35.72 | 1173 | Node5 | 31.01 |
| 511 | M_glareolus_v2 | 712.62 | 392 | Node4 | 38.37 | 1184 | Node5 | 55.95 |
| 640 | M_glareolus_v2 | 1766.18 | 393 | Node4 | 193.92 | 1196 | Node5 | 10.9 |

**Table S2:** Summary results from the population program in Stacks using all filtered, thinned SNPs. Pop ID: population ID; Num_Indv: average number of individuals at each SNP; P: mean frequency of the most common allele at each SNP; Obs_Het: mean observed heterozygosity; Obs_Hom: mean observed homozygosity; Exp_Het: mean expected heterozygosity; Exp_Hom: mean expected homozygosity; Pi: mean nucleotide diversity (π); Fis: mean inbreeding coefficient (F_IS_); StdErr: standard error; dab: Dąbrowica; TEL: Teleśnica; jul: Julianka; bial: Białystok; NIEP: Niepołomice; aug: Augustów; ol: Olsztyn; elk: Ełk; ziel: Zielonka; socz: Soczewka; zmig: Zmigród.

| Pop ID | Num_Indv | StdErr | P | StdErr | Obs_Het | StdErr | Obs_Hom | StdErr | Exp_Het | StdErr | Exp_Hom | StdErr | Pi | StdErr | Fis | StdErr |
| --- | --- | --- | --- | --- | --- | --- | --- | --- | --- | --- | --- | --- | --- | --- | --- | --- |
| dab | 17.62 | 0.00 | 0.83 | 0.00 | 0.24 | 0.00 | 0.76 | 0.00 | 0.24 | 0.00 | 0.76 | 0.00 | 0.24 | 0.00 | 0.02 | 0.00 |
| TEL | 12.28 | 0.00 | 0.83 | 0.00 | 0.24 | 0.00 | 0.76 | 0.00 | 0.24 | 0.00 | 0.76 | 0.00 | 0.25 | 0.00 | 0.03 | 0.00 |
| jul | 8.92 | 0.00 | 0.84 | 0.00 | 0.21 | 0.00 | 0.79 | 0.00 | 0.22 | 0.00 | 0.78 | 0.00 | 0.23 | 0.00 | 0.05 | 0.00 |
| bial | 10.31 | 0.00 | 0.84 | 0.00 | 0.24 | 0.00 | 0.76 | 0.00 | 0.23 | 0.00 | 0.77 | 0.00 | 0.24 | 0.00 | 0.01 | 0.00 |
| NIEP | 13.42 | 0.00 | 0.83 | 0.00 | 0.23 | 0.00 | 0.77 | 0.00 | 0.23 | 0.00 | 0.77 | 0.00 | 0.24 | 0.00 | 0.04 | 0.00 |
| brok | 10.81 | 0.00 | 0.84 | 0.00 | 0.24 | 0.00 | 0.76 | 0.00 | 0.23 | 0.00 | 0.77 | 0.00 | 0.24 | 0.00 | 0.02 | 0.00 |
| aug | 6.98 | 0.00 | 0.84 | 0.00 | 0.23 | 0.00 | 0.77 | 0.00 | 0.22 | 0.00 | 0.78 | 0.00 | 0.24 | 0.00 | 0.02 | 0.00 |
| ol | 10.99 | 0.00 | 0.84 | 0.00 | 0.23 | 0.00 | 0.77 | 0.00 | 0.23 | 0.00 | 0.77 | 0.00 | 0.24 | 0.00 | 0.03 | 0.00 |
| elk | 13.52 | 0.00 | 0.83 | 0.00 | 0.23 | 0.00 | 0.77 | 0.00 | 0.23 | 0.00 | 0.77 | 0.00 | 0.24 | 0.00 | 0.03 | 0.00 |
| ziel | 14.85 | 0.00 | 0.83 | 0.00 | 0.23 | 0.00 | 0.77 | 0.00 | 0.23 | 0.00 | 0.77 | 0.00 | 0.24 | 0.00 | 0.05 | 0.00 |
| gj | 5.69 | 0.00 | 0.84 | 0.00 | 0.22 | 0.00 | 0.78 | 0.00 | 0.21 | 0.00 | 0.79 | 0.00 | 0.23 | 0.00 | 0.03 | 0.00 |
| socz | 6.54 | 0.00 | 0.84 | 0.00 | 0.22 | 0.00 | 0.78 | 0.00 | 0.22 | 0.00 | 0.78 | 0.00 | 0.24 | 0.00 | 0.04 | 0.00 |
| zmig | 5.66 | 0.00 | 0.84 | 0.00 | 0.23 | 0.00 | 0.77 | 0.00 | 0.22 | 0.00 | 0.78 | 0.00 | 0.24 | 0.00 | 0.02 | 0.00 |

**Table S3:** Populations sampled for CFH and RAD sequencing. # samples = number of remaining samples per population after filtering of the RAD-seq data.

| Population | Longitude | Latitude | # samples |
| --- | --- | --- | --- |
| Augustów | N53.796200 | E23.150766 | 8 |
| Białystok | N53.279194 | E23.373034 | 11 |
| Brok | N52.700283 | E21.918842 | 12 |
| Dąbrowica | N50.475753 | E22.359139 | 22 |
| Ełk | N53.801549 | E22.399108 | 15 |
| Goly Jon | N53.693694 | E18.151994 | 6 |
| Julianka | N50.773350 | E19.465696 | 11 |
| Niepołomice | N50.00000 | E20.200000 | 14 |
| Olsztyn | N53.516669 | E20.624690 | 12 |
| Soczewka | N52.534904 | E19.598152 | 7 |
| Teleśnica | N49.220000 | E22.320000 | 14 |
| Zielonka | N52.582626 | E17.152514 | 16 |
| Zmigród | N51.508057 | E17.056494 | 6 |

**Table S4:** IDs and populations of origin of individuals sampled for full-length CFH sequencing.

| Bank vole ID | Population |
| --- | --- |
| R122 | Brok |
| R129 | Brok |
| R151 | Olsztyn |
| R152 | Olsztyn |
| R286 | Julianka |
| R304 | Żmigród |
| R306 | Żmigród |
| 009_22 | Ełk |
| 017_22 | Ełk |
| R149 | Olsztyn |
| R150 | Olsztyn |
| R262 | Goły Jon |
| R263 | Goły Jon |
| R264 | Goły Jon |
| R287 | Julianka |
| R288 | Julianka |
| R289 | Julianka |
| R303 | Żmigród |
| R305 | Żmigród |

**Table S5:** IDs and populations of the individuals sampled to generate a database of expressed CFH.

| Bank vole ID | Population ID | individuals/population |
| --- | --- | --- |
| R_4 | Brok | 9 |
| R_6 | Brok |  |
| R_2 | Brok |  |
| R_54 | Brok |  |
| R_55 | Brok |  |
| R_57 | Brok |  |
| R_58 | Brok |  |
| R_59 | Brok |  |
| R_60 | Brok |  |
| R_13 | Ełk | 12 |
| R_14 | Ełk |  |
| R_16 | Ełk |  |
| R_17 | Ełk |  |
| R_18 | Ełk |  |
| R_16 | Ełk |  |
| R_33 | Ełk |  |
| R_34 | Ełk |  |
| R_36 | Ełk |  |
| R_37 | Ełk |  |
| R_39 | Ełk |  |
| R_40 | Ełk |  |
| R_19 | Olsztyn | 11 |
| R_20 | Olsztyn |  |
| R_21 | Olsztyn |  |
| R_22 | Olsztyn |  |
| R_23 | Olsztyn |  |
| R_24 | Olsztyn |  |
| R_25 | Olsztyn |  |
| R_26 | Olsztyn |  |
| R_27 | Olsztyn |  |
| R_28 | Olsztyn |  |
| R_29 | Olsztyn |  |
| R_41 | Augustów | 6 |
| R_42 | Augustów |  |
| R_43 | Augustów |  |
| R_44 | Augustów |  |
| R_45 | Augustów |  |
| R_46 | Augustów |  |
| R_52 | Białystok | 2 |
| R_53 | Białystok |  |
| R_61 | Zielonka | 7 |
| R_62 | Zielonka |  |
| R_63 | Zielonka |  |
| R_66 | Zielonka |  |
| R_67 | Zielonka |  |
| R_68 | Zielonka |  |
| R_69 | Zielonka |  |
| R_8 | Żebra Żubra | 1 |
| R_9 | Dubeczno | 1 |
| R_11 | Dąbrowica | 1 |

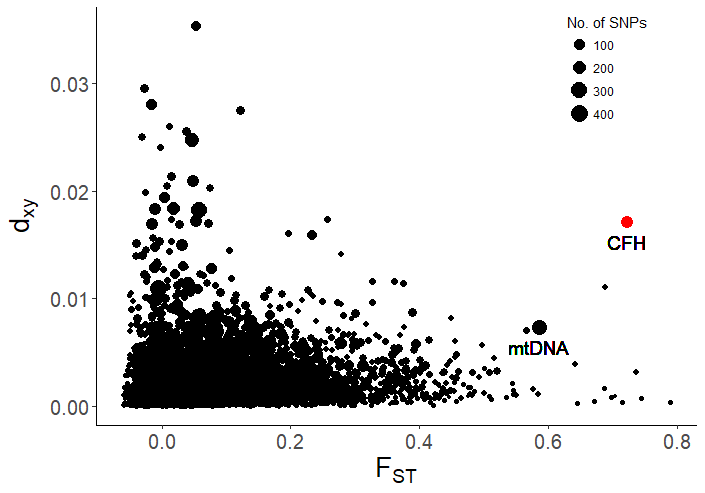

**Figure S1: CFH is among the most differentiated and diverged genes between Eastern and Western bank vole populations.**
Differentiation and divergence between Eastern (Białowieża) and Western (Włocławek) populations of bank voles. The plot is based on transcriptomic data from Niedziałkowska et al. (2023) which represent 10 individuals from Białowieża and 8 individuals from Włocławek populations. Each dot represents a single transcript. The transcript annotated as CFH is shown in red.

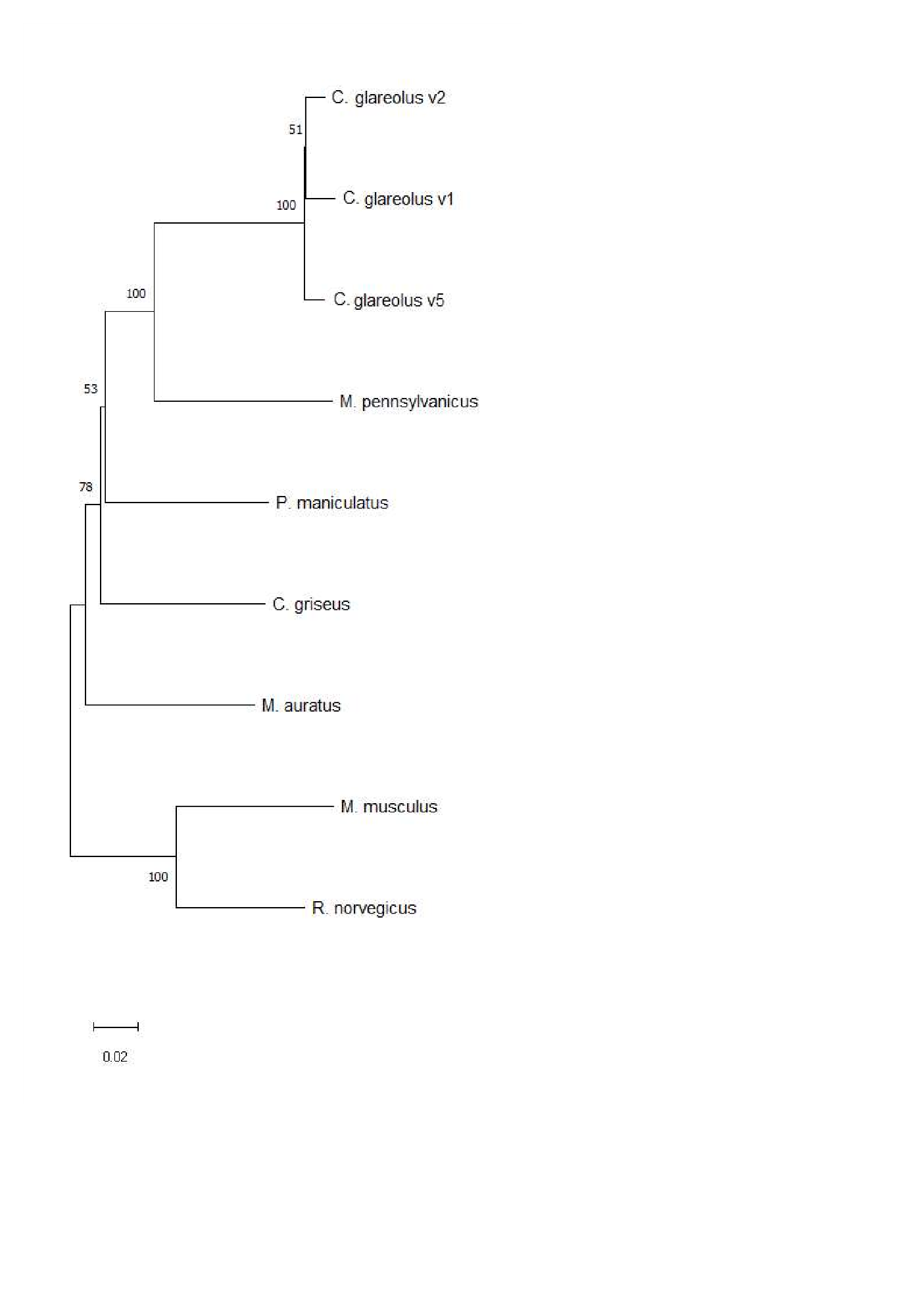

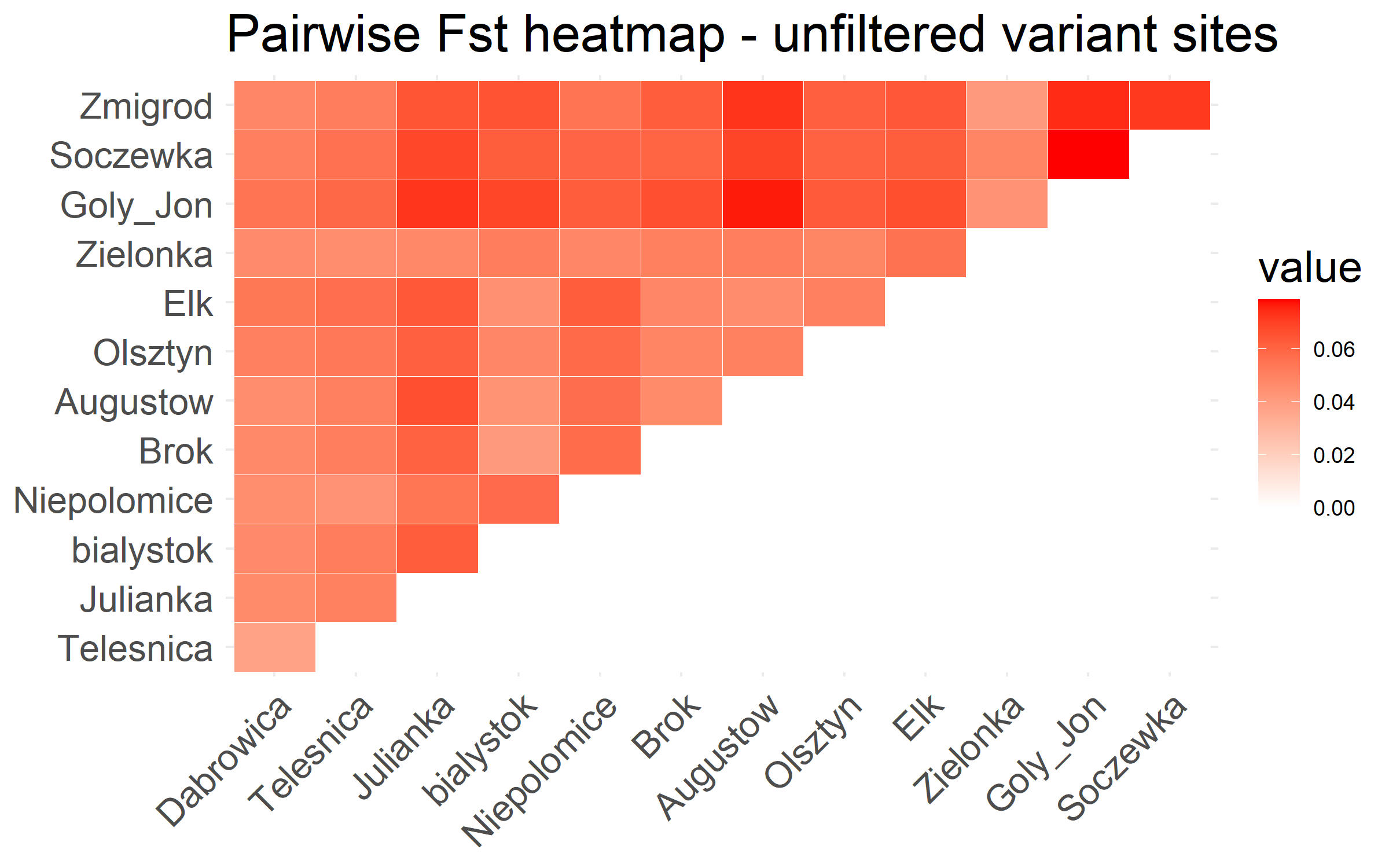

**Figure S3: Populations show reduced genomic differentiation.**
Pairwise F_ST_ heatmap generated using the results from the population program in Stacks using all filtered, thinned SNPs.

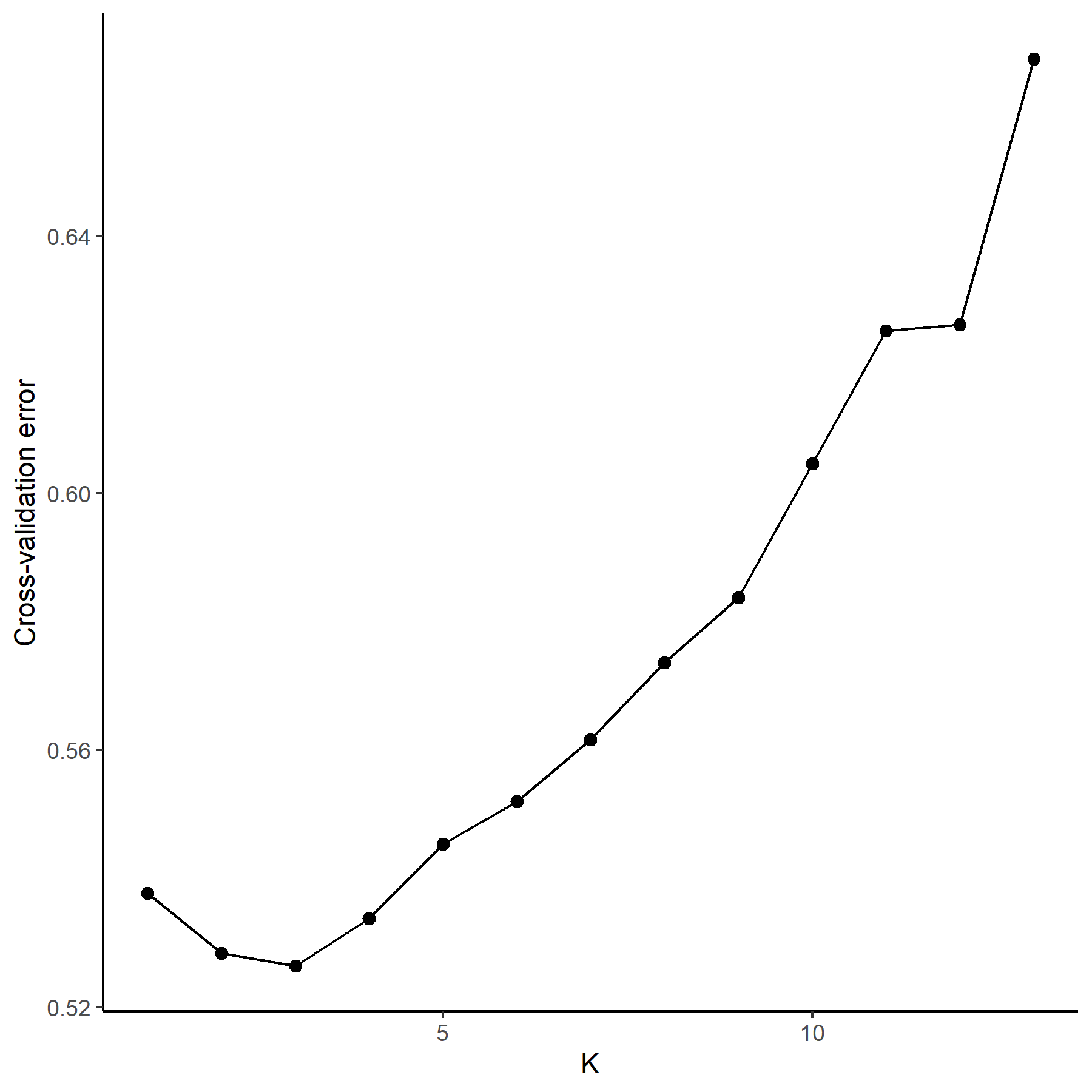

**Figure S4: Populations cluster in three main groups.**
X-axis: number of groups (K); y-axis: cross-validation error.

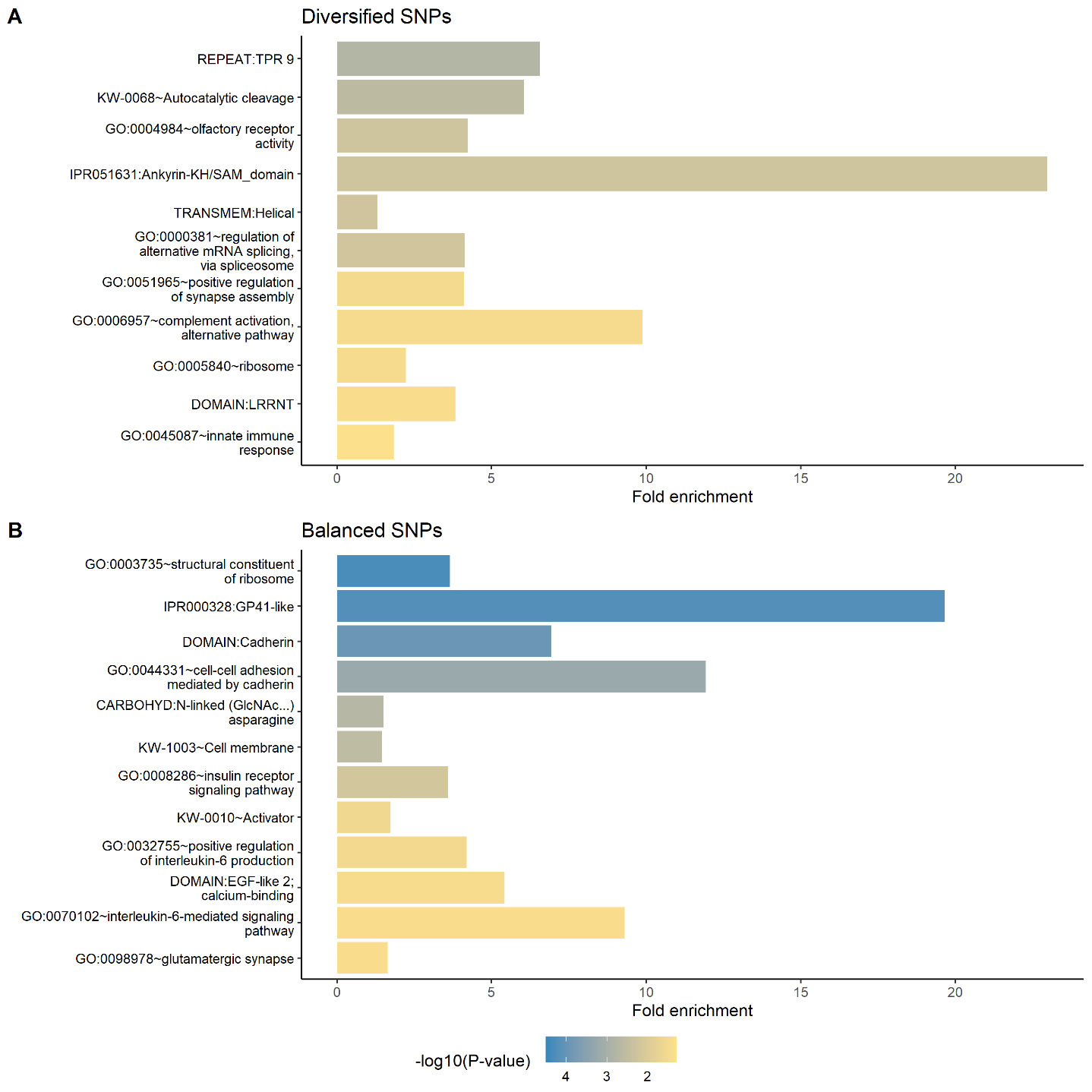

**Figure S5: SNPs under positive selection are enriched for terms related to alternative complement activation and innate immunity, while those under balancing selection for terms related to Interleukin 6 pathways.**
Enrichment tests using the overlapping or closest genes to the SNPs under diversifying (a) or balancing (b) selection. X-axis: fold enrichment; colors of the bars: -log_10_ of p-values. Enriched terms are clustered based on function; one representative term for each cluster is shown.

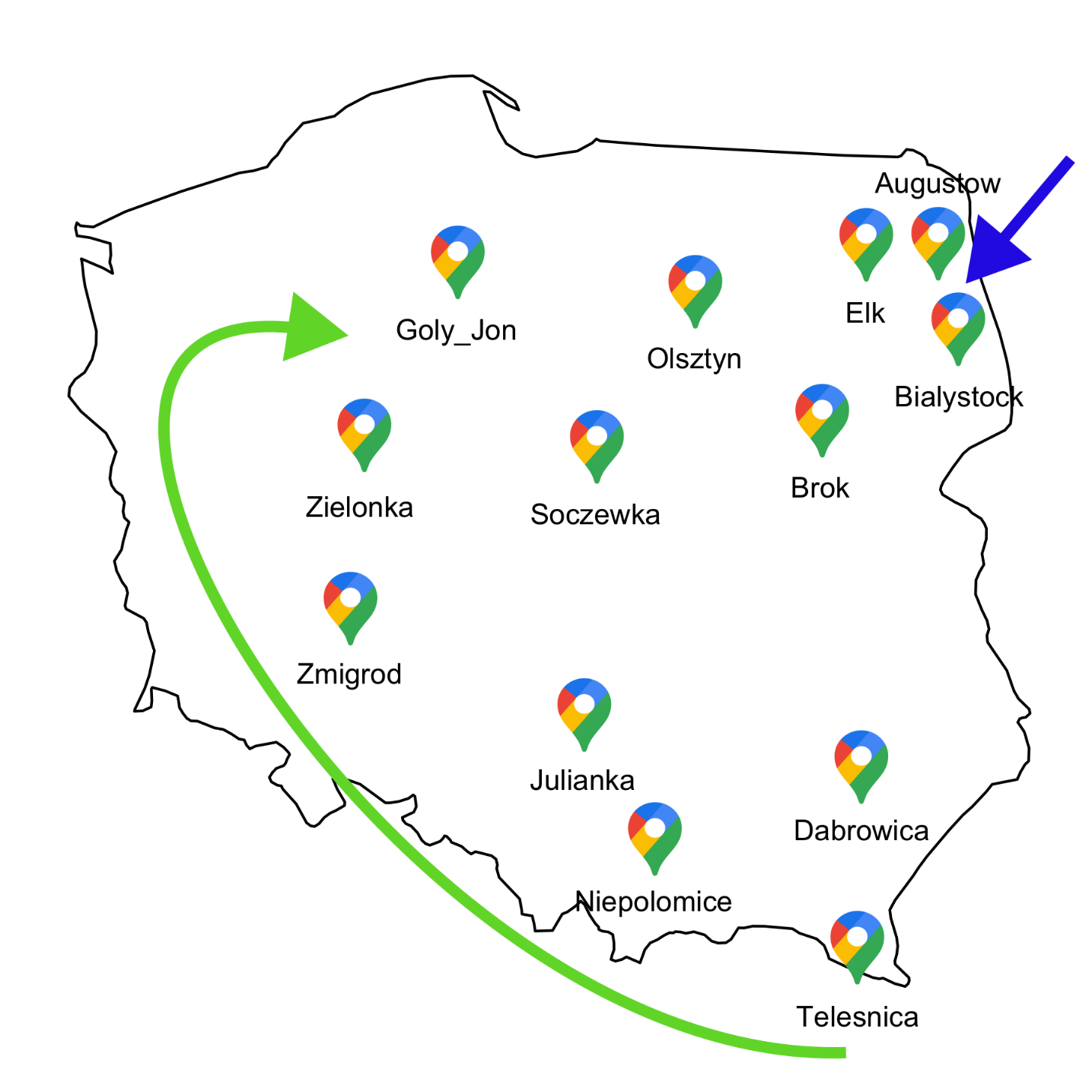

**Figure S6: The sampled populations span the postglacial colonization areas and the contact zone of the Carpathian and the Eastern bank vole lineages.**
Map of Poland showing the location of the populations sampled for CFH and RAD sequencing. Green arrow: postglacial colonization area of the Carpathian clade; blue arrow: postglacial colonization area of the Eastern clade.

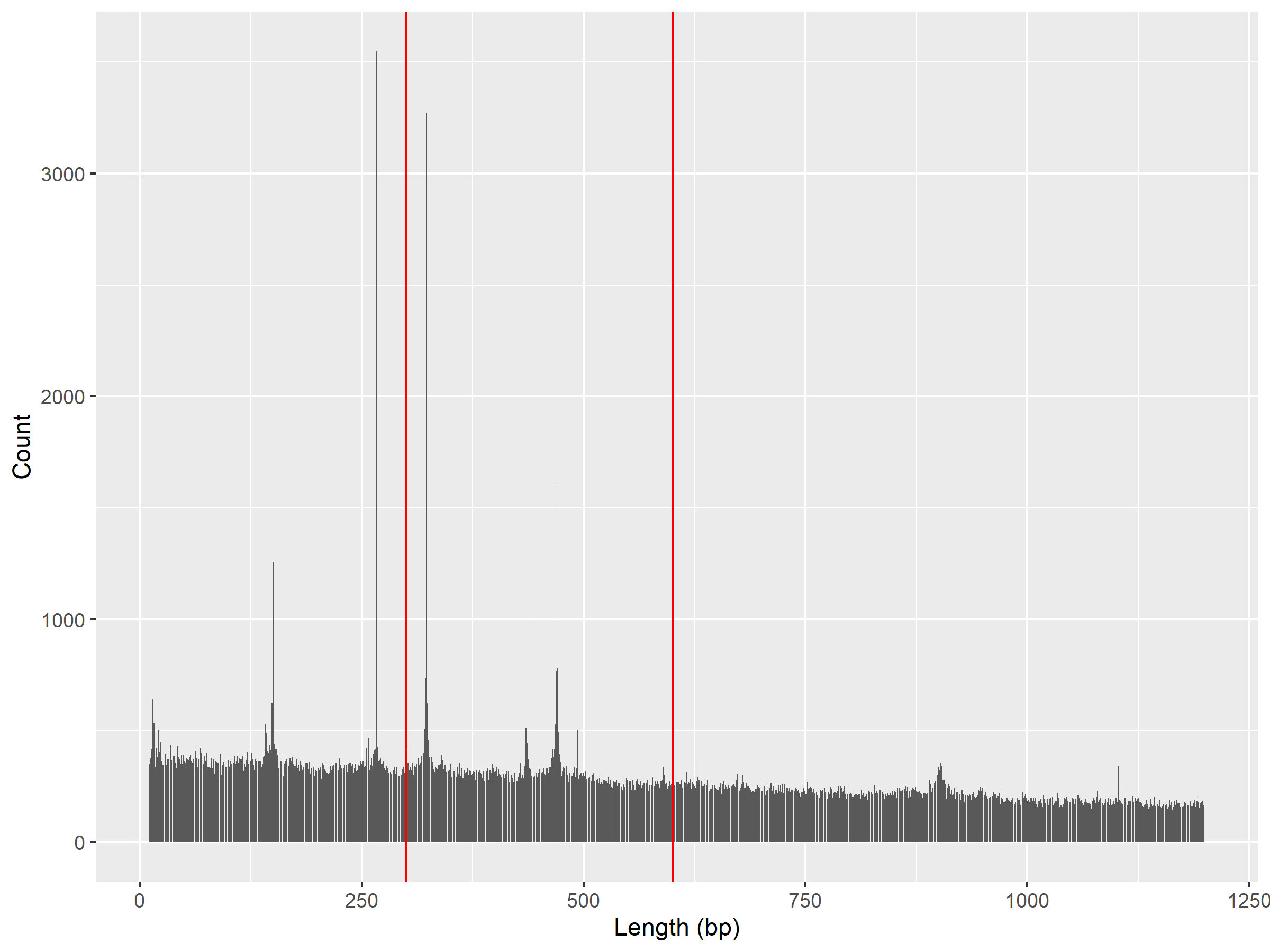

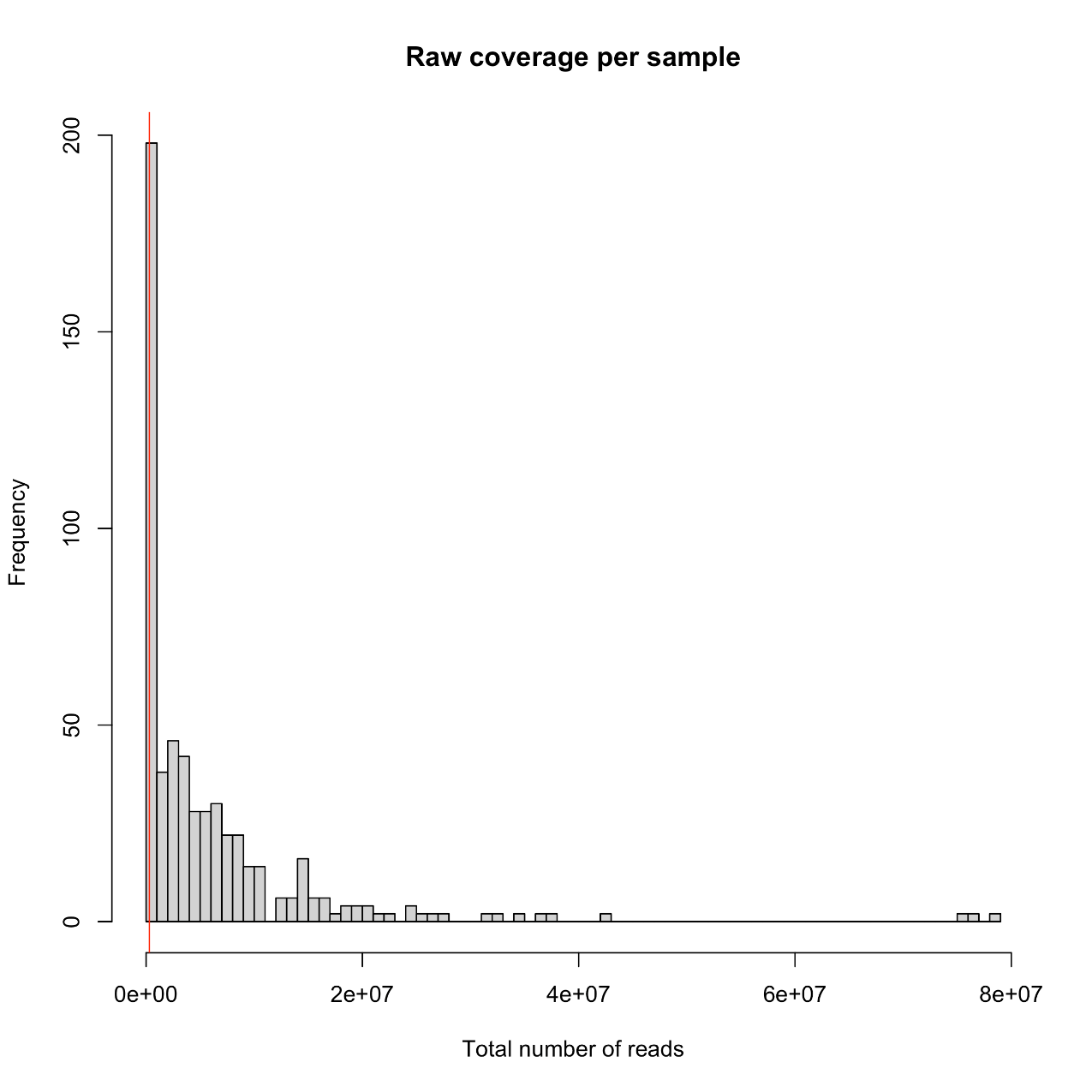

**Figure S8: Histogram of the raw coverages per sample.**
The red bar shows 10% of the median of the raw coverage across all samples (= 302,396.5 reads). The samples below this threshold were discarded. Coverages were calculated for each forward and reverse file of each sample.

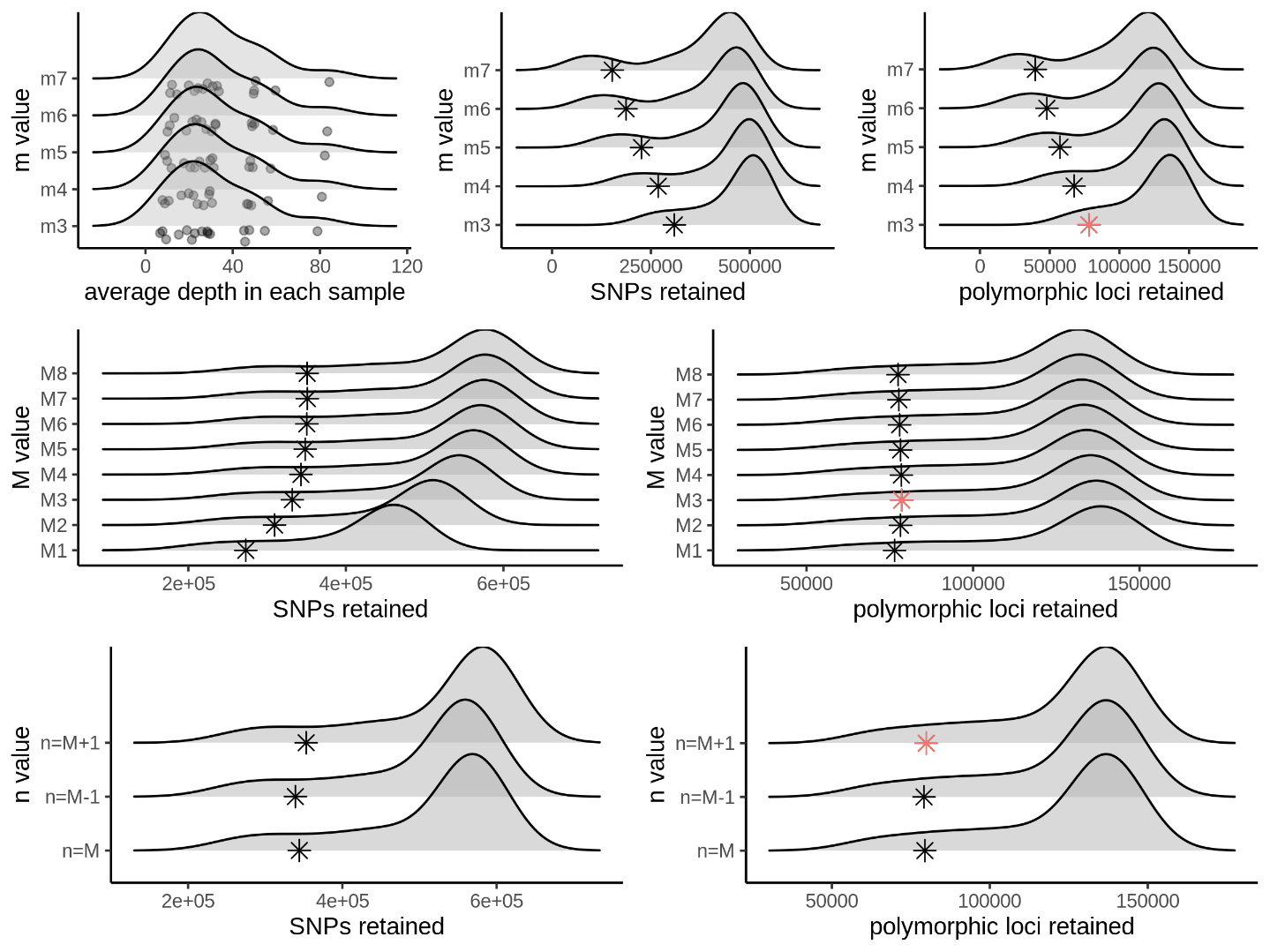

**Figure S9: Stacks parameter optimizations.**
(Top) *m* parameter optimization shown in density plots of the average depth in each sample (left), number of Single Nucleotide Polymorphisms (SNPs) retained (middle), and number of polymorphic sites retained (right) at each value of *m* = 3-7. (Middle) *M* parameter optimization shown in density plots of the number of SNPs retained (left) and the number of polymorphic loci retained (right) for each value of *M* = 1-8. (Bottom) *n* parameter optimization shown in density plots of the number of SNPs retained (left) and the number of polymorphic loci retained (right) for each value of *n*: *n* = *M* = 3; *n* = *M* – 1 = 2; *n* = *M* + 1 = 4. The red stars in the rightmost plots show the optimal value for each parameter.

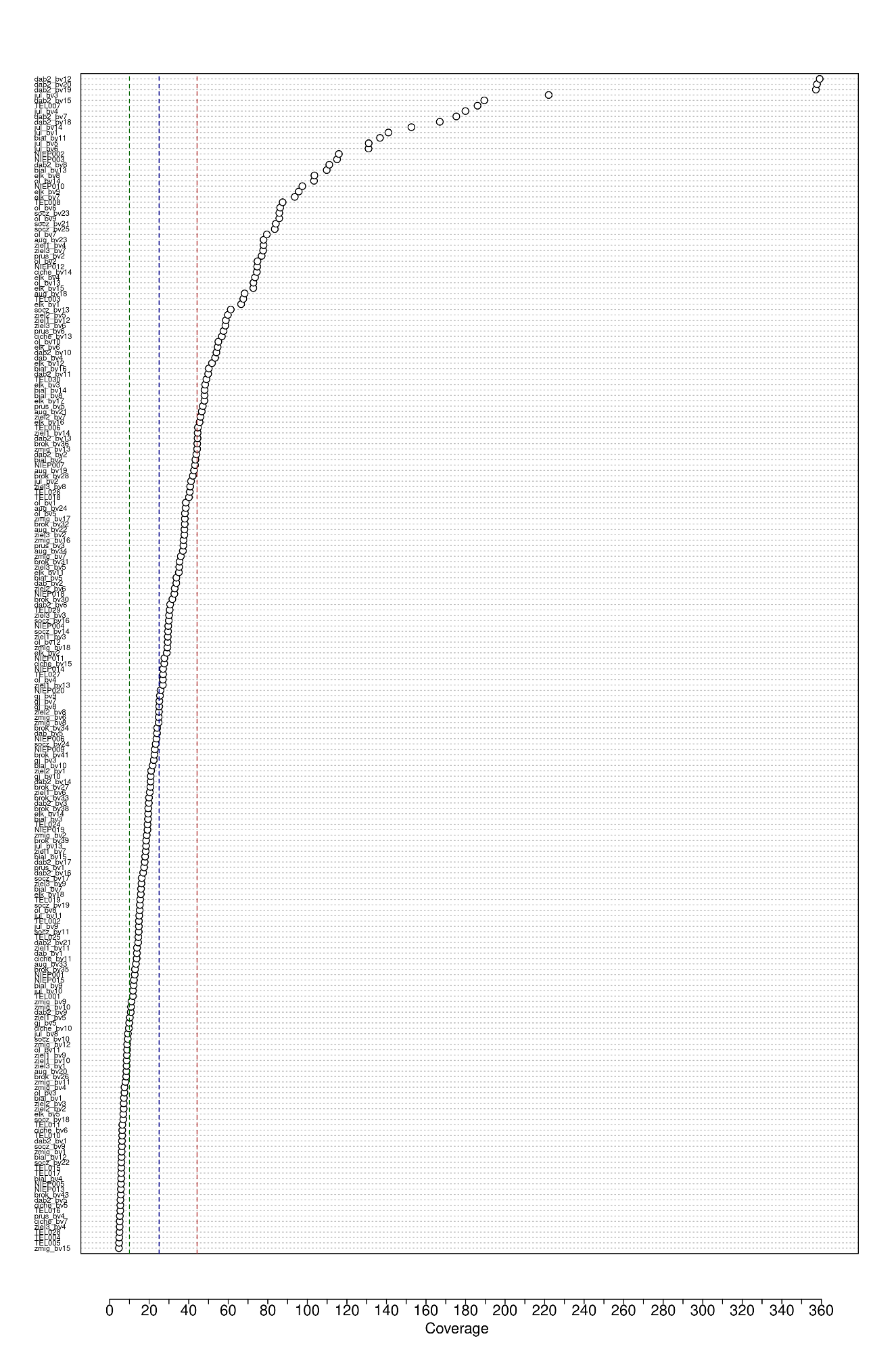

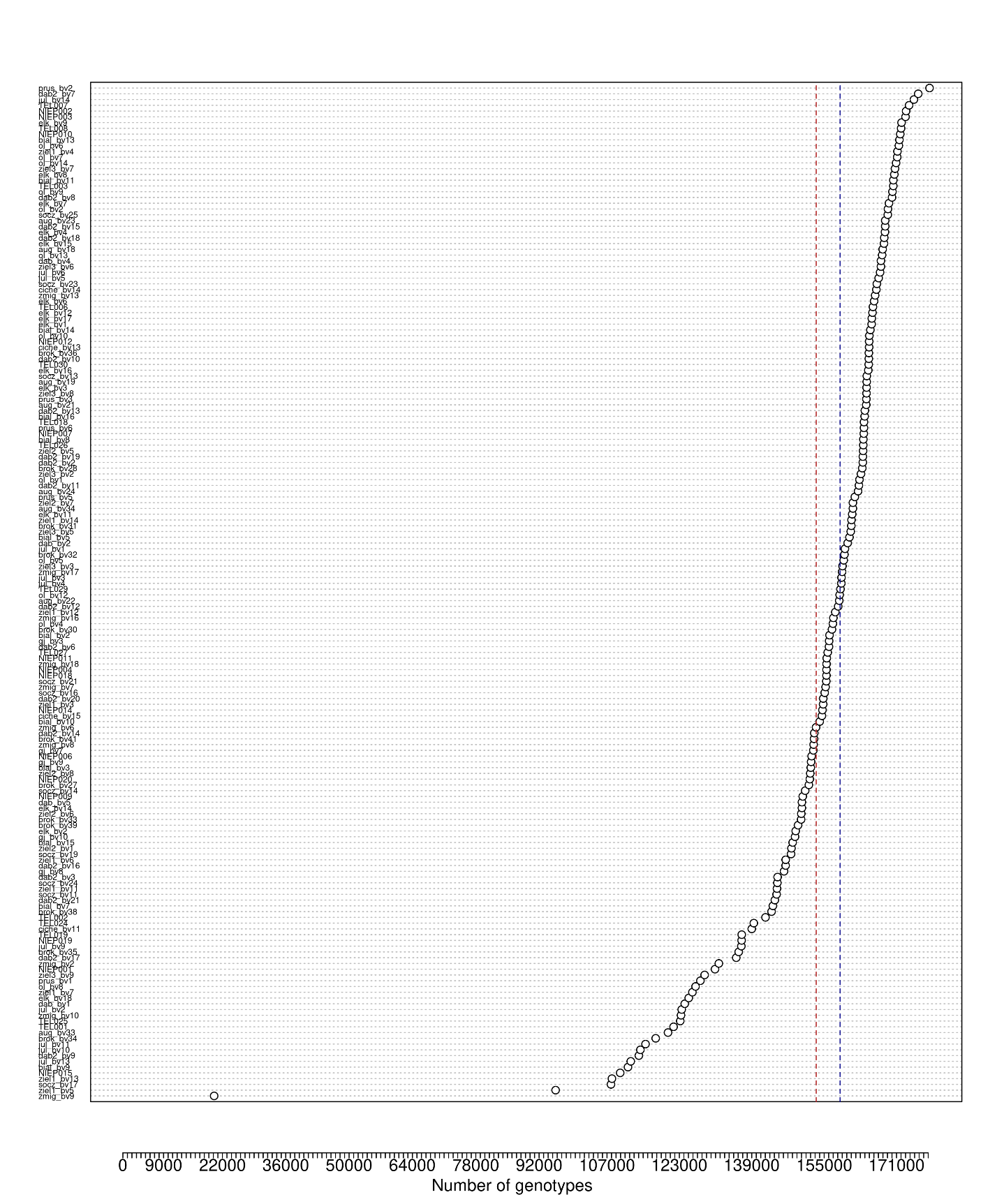

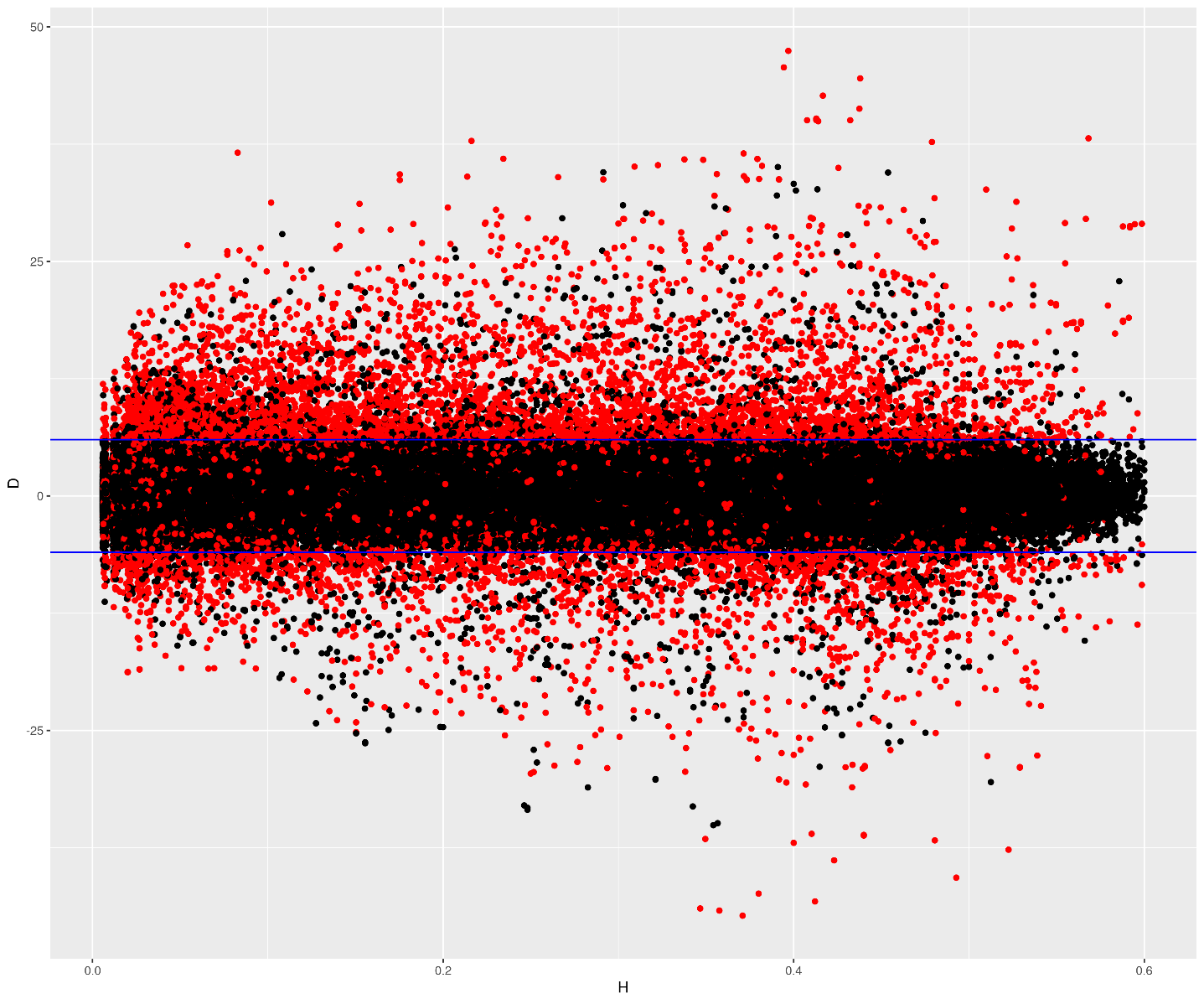

**Figure S11: Paralogous loci check.**
The graph shows the results of ‘HDPLOT’. X-axis = expected proportion of heterozygous individuals; y-axis = deviation from the 1:1 allelic ratio for heterozygotes. Blue lines = chosen cut-off for the read ratio deviation (|D| = 6). Black dots = SNPs in non-paralogous loci; red dots = SNPs in paralogous loci identified by both ‘HDPlot’ and VSEARCH (3,600 loci, discarded from further analyses). Loci with at least one SNP with |D| >6 in ‘HDPlot’ were considered paralogous; that is why some SNPs were discarded even though they had |D| <6.

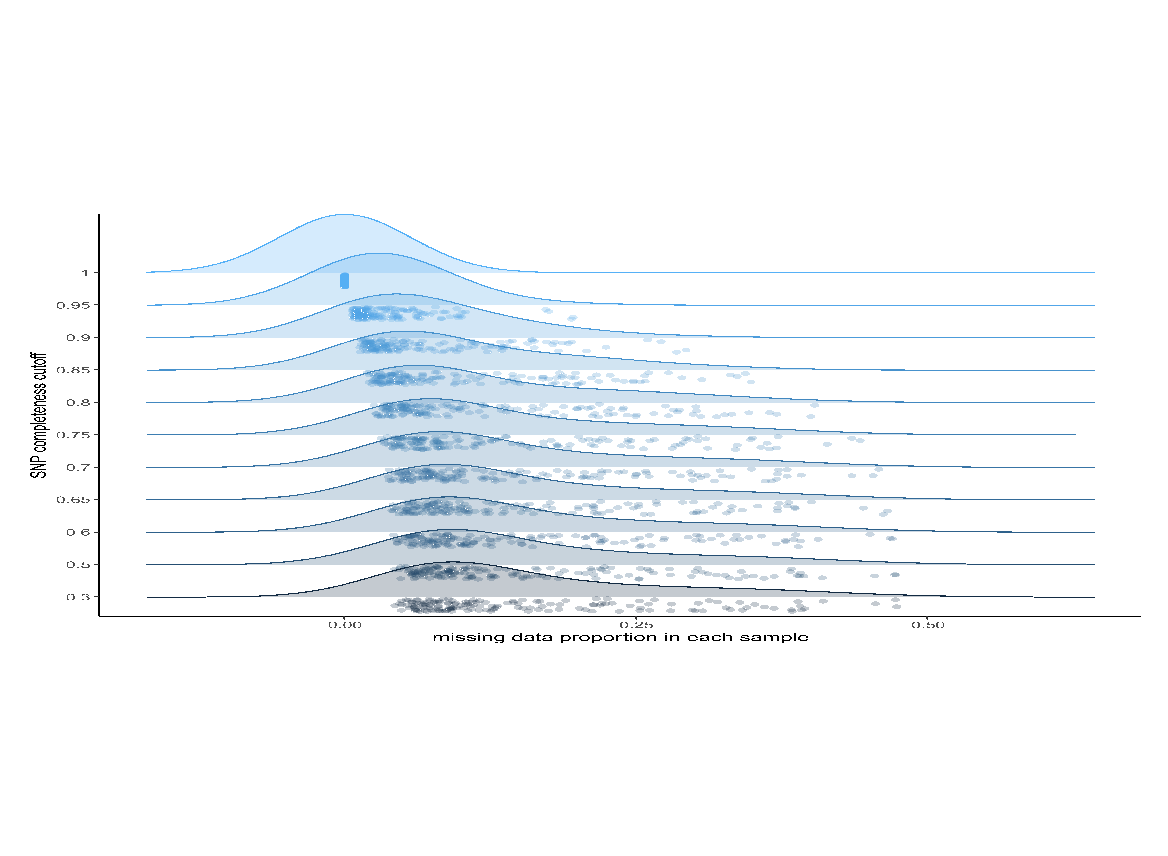

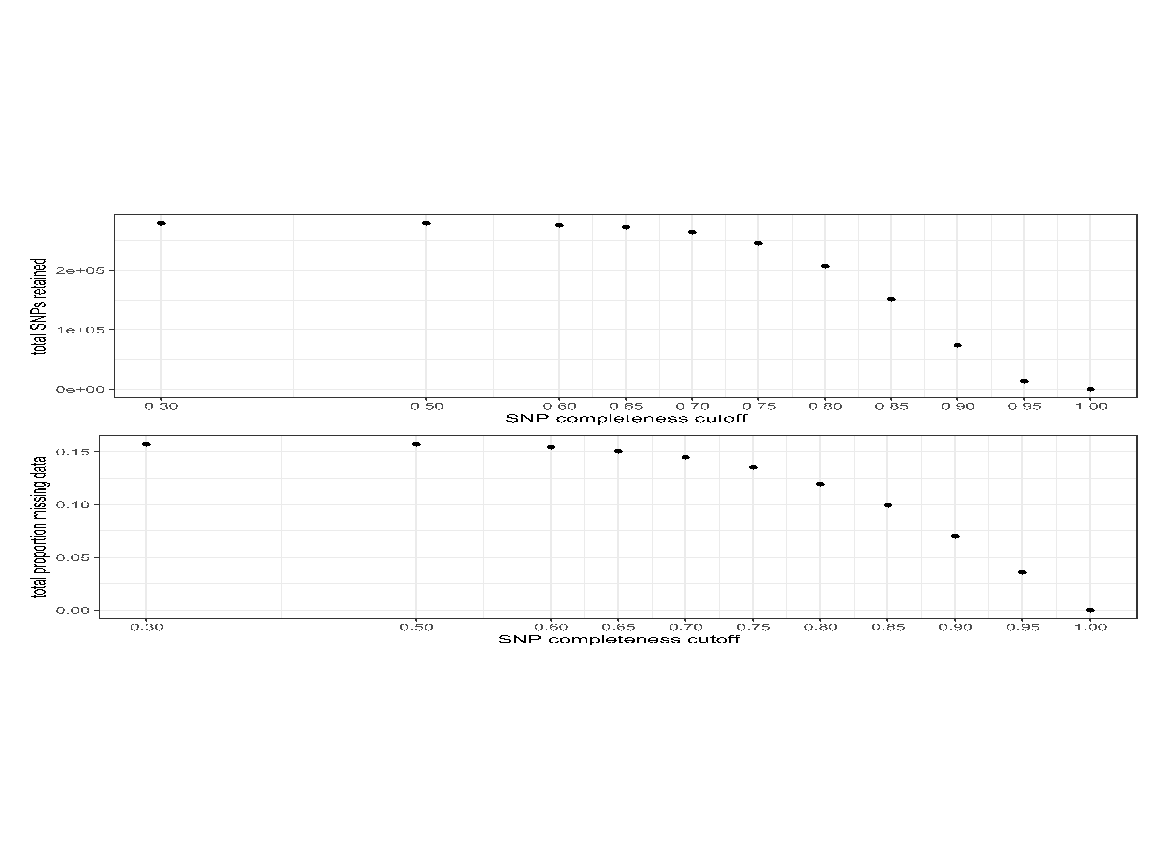

**Figure S12: SNP completeness cutoff effects on missing data and number of SNPs.**
(A) Density plot showing the proportion of missing data per sample for each SNP completeness (the opposite of missing data per SNP) cutoff (0.3-1). Dots are samples. (B) Total number of SNPs retained and total proportion of missing data by SNP completeness cutoff. The chosen cutoff (0.8) represents a tradeoff between reducing the differences in missing data between samples and retaining the most SNPs.
